# TWIST1 interacts with adherens junction proteins during neural tube development and regulates fate transition in cranial neural crest cells

**DOI:** 10.1101/2021.08.22.457283

**Authors:** Jessica W. Bertol, Shelby Johnston, Rabia Ahmed, Victoria K. Xie, Kelsea M. Hubka, Lissette Cruz, Larissa Nitschke, Marta Stetsiv, Jeremy P. Goering, Paul Nistor, Sally Lowell, Hanne Hoskens, Peter Claes, Seth M. Weinberg, Irfan Saadi, Mary C. Farach-Carson, Walid D. Fakhouri

## Abstract

Cell fate determination is a necessary and tightly regulated process for producing different cell types and structures during development. Cranial neural crest cells (CNCCs) are unique to vertebrate embryos and emerge from the neural fold borders into multiple cell lineages that differentiate into bone, cartilage, neurons, and glial cells. We previously reported that *Irf6* genetically interacts with *Twist1* during CNCC-derived tissue formation. Here, we investigated the mechanistic role of *Twist1* and *Irf6* at early stages of craniofacial development. Our data indicates that TWIST1 interacts with α/β/γ-CATENINS during neural tube closure, and *Irf6* is involved in the structural integrity of the neural tube. *Twist1* suppresses *Irf6* and other epithelial genes in CNCCs during epithelial-to-mesenchymal transition (EMT) process and cell migration. Conversely, a loss of *Twist1* leads to a sustained expression of epithelial and cell adhesion markers in migratory CNCCs. Disruption of TWIST1 phosphorylation *in vivo* leads to epidermal blebbing, edema, neural tube defects, and CNCC-derived structural abnormalities. Altogether, this study describes an uncharacterized function of *Twist1* and *Irf6* in the neural tube and CNCCs and provides new target genes of *Twist1* involved in cytoskeletal remodeling. Furthermore, the association between DNA variations within TWIST1 putative enhancers and human facial morphology is also investigated.

**SUMMARY STATEMENT:** This study uncovers a new function of *Twist1* in neural tube development and epithelial-to-mesenchymal transition in cranial neural crest cells. Data further shows that *Twist1*-interacting *Irf6* is involved in regulating neural tube integrity.

## INTRODUCTION

Craniofacial morphogenesis requires precise interaction between multiple genes and gene-environments that are essential for cell fate induction, specification, and differentiation (Chai and Maxon Jr., 2006; Murray et al., 2007; Nomura et al., 1998; Padmanabhan and Ahmed, 1997; Peyrard-Janvid et al., 2014; Sato et al., 2008). Disruption of the precisely organized morphogenesis leads to craniofacial disorders, which are the second most common congenital birth defects that occur either as part of syndromes or as isolated phenotypes (Hoyert et al., 2006; Joshi et al., 2014; Parker et al., 2010). Neuroectoderm, which becomes the neural plate, is induced by signals that emerge from the notochord and mesoderm. The junctions between non-neural and neural ectoderm are called the neural plate borders, which form distinct domains through inductive interactions with the adjacent cells and the underlying mesoderm. Signaling molecules, including fibroblast growth factors (FGFs), wingless-related integration sites (WNTs), and bone morphogenetic proteins (BMPs), induce CNCCs and placodal progenitor cells by activating downstream transcription factors (TFs) for cell specification (Groves and LaBonne, 2014; Litingtung and Chiang, 2000; Soo et al., 2002; Millonig et al., 2000). The CNCCs emerge from the neural fold borders of the mid- and hindbrain regions and migrate as unipotent, bipotent, and multipotent mesenchymal cells towards the pouches of the frontonasal and pharyngeal arches (Achilleos and Trainor, 2012). CNCCs give rise to the bone, cartilage, neuron, and glial cells of the frontal and orofacial skeletal and peripheral nervous systems (Groves and LaBonne, 2014; He et al., 2014; Le Douarin et al., 2004; Noden and Trainor, 2005).

*Twist1* encodes a transcription factor that belongs to the basic helix-loop-helix (bHLH) B-family. It was first identified in *Drosophila melanogaster* where its deletion resulted in disruption of mesodermal cell fate and ventral furrow formation at the gastrulation stage (Lu et al., 2011; Nusslein-Volhard et al., 1984; Simpson, 1983). In vertebrates, *Twist1* is expressed first at the gastrulation stage in the primitive streak and then in the adjacent mesodermal cells (Fuchtbauer, 1995; Stoetzel et al., 1995). At the neurulation stage, *Twist1* is detected in the paraxial and lateral mesoderm and the migratory CNCCs (Fuchtbauer, 1995; Soo et al., 2002). *Twist1* expression is maintained in the cranial neural crest-derived mesenchyme of the frontonasal and pharyngeal processes (Bildsoe et al., 2009; Fuchtbauer, 1995). In humans, mutations in *TWIST1* cause Saethre-Chotzen syndrome, characterized by craniosynostosis and cleft palate, Sweeney-Cox syndrome, and Robinow-Sorauf syndrome (Kunz and Fritz, 1999; Takenouchi et al., 2018; Seto et al., 2007). In mice, *Twist1* is crucial for neural tube closure and CNCC-derived craniofacial bone and cartilage formation as reported in the phenotypes of *Twist1* null and conditional knockout (CKO) in CNCCs (Chen et al., 2007; Bildsoe et al., 2013; Zhang et al., 2012). TWIST1 is shown to promote cell survival and proliferation of migratory CNCCs during craniofacial development (Bildsoe et al., 2013). Although both processes involve gaining potency and motility, the *in vivo* molecular function of TWIST1 as a master regulator of epithelial-to-mesenchymal transition (EMT) in CNCCs is still unknown (Fan, et al., 2021).

TWIST1 is a phosphoprotein with multiple residues that are phosphorylated by several kinases according to numerous cancer cell line studies (Lu et al., 2011; Vichalkovski et al., 2010). TWIST1 phosphorylation is important for its homo- and heterodimerization with other transcription factors to regulate multiple cellular activities (Hong et al., 2011; Lu et al., 2011; Vichalkovski et al., 2010). Bioinformatic analysis identified 9 putative phosphorylation sites of TWIST1 (Hong et al., 2011). TWIST1 phosphorylation at serine 18 (S18) and S20 increases tumor cell motility in squamous cell carcinoma of the head and neck (SCCHN) cancer cells (Su et al., 2011). Furthermore, phosphorylation of S68 by MAPK is required for protein stability and breast cancer cell invasiveness (Hong et al., 2011), whereas threonine (T) 125 and S127 are crucial phosphorylation sites for protein dimerization found in the bHLH domain. Human genetic studies showed that disruption of phospho-sites in the bHLH region causes autosomal dominant craniosynostosis (Firulli et al., 2005; Seto et al., 2007). In mice, overexpression of *Twist1^T125D/S127D^* phosphomimetic alleles (equivalent to T121/S123 in human) in cardiomyocytes resulted in disrupted cardiac cell remodeling and heart formation of transient transgenic embryos leading to sudden death. In contrast, overexpression of phospho-incompetent *Twist1^T125A/S127A^* alleles did not show heart pathology in transient transgenic embryos (Lu et al., 2011). TWIST1 proteins bind to DNA as homodimers and can interact with proteins like TCF3 and other chromatin remodeling proteins to guide CNCC migration (Fan et al., 2020). TCF3 can usually form a complex with LEF1 and β-CATENIN to regulate target gene expression (Fakhouri et al., 2012).

Our prior work demonstrated that compound heterozygous mice (*Irf6^+/−^; Twist1^+/−^*) have craniofacial abnormalities, namely mandibular agnathia, fused maxilla, cleft palate, and holoprosencephaly (Fakhouri et al., 2017). Unlike other IRF family members, *Interferon Regulatory Factor 6* (*IRF6*) is a broadly expressed transcription factor in the ectoderm and oral epithelium during embryogenesis. In mice, *Irf6* promotes cell differentiation of proliferative epithelial cells, but its function in the neural tube and neural crest cell development is still unknown (Ingraham et al., 2006; Kousa et al., 2019; Richardson et al. 2005). We recently showed that overexpression of *Irf6* in ectoderm by K14-enhancer leads to exencephaly and failure of peritoneal skin development. Furthermore, the compound heterozygous mice for *Irf6* and *Ap2*α have skin protrusions and deformities in the ectoderm along the neural tube (Kousa et al., 2019).

In the current study, we sought to characterize the spatiotemporal expression of *Twist1* and *Irf6* during early stages of neural tube formation and their role in CNCC development. We generated multiple stable *Twist1* phospho-incompetent mouse lines to identify the importance of TWIST1 phosphorylation in regulating neural tube and CNCC-derived craniofacial structures. We investigated the function of *Irf6* in regulating the integrity of the neural tube and its relation to restricting AP2α expression to the junction with non-neural ectoderm. Our data shows that TWIST1 expression pattern overlaps with adherens and tight junction proteins in neural tube and TWIST1 interacts with α/β/ɣ-CATENINS during neural tube closure. We also show that *Twist1* is crucial to promote cell fate transition of neuroepithelial cells to become migratory mesenchymal cells and loss of *Twist1* in CNCCs causes ectopic expression of IRF6 and E-CADHERIN in delaminated epithelial-like neural crest cells. We demonstrate that three highly conserved phospho-residues in TWIST1 are crucial for its *in vivo* function in the brain and CNCC-derived craniofacial structures. Finally, our data shows that TWIST1 regulates *Specc1l* expression during CNCC development by binding to its putative regulatory elements to possibly control cell adhesion and cytoskeletal remodeling in migratory CNCCs.

## RESULTS

### Spatiotemporal expression of TWIST1 and IRF6 during neural tube development

To determine the role of *Twist1* and *Irf6* in neural tube formation, we investigated the spatiotemporal expression at the early stages of neural tube formation in wild type (WT) embryos. We performed immunofluorescent (IF) staining and *in situ* hybridization in WT embryos from E8.5 to E9.5. At E8.5, IRF6 is highly expressed throughout the neural plate and in a few mesodermal cells (Fig.1A, A’), while TWIST1 is unexpectedly detected in spot-like structures at the apical side of the neural plate (Fig 1A, A’ white arrow). TWIST1 was also expressed in a few cells at the neural plate borders (Fig. 1SA, A’, D, D’). The nuclear expression of TWIST1 was observed in the mesodermal cells adjacent to the neural plate (Fig. 1A’, B, B’, green arrow). At E9.0, most cells in neural folds express IRF6, while TWIST1 is similarly expressed at the apical side of the neural folds. TWIST1 expression showed an overlap with IRF6 at the apical side and in some cells at the dorsal edges of the neural folds (Fig. 1E, E’, white arrow). To understand the significance of the apical expression in neural folds, we investigated whether the TWIST1 expression pattern overlaps with adherens and tight junction proteins. TWIST1 apical expression in the dorsal neuroectodermal cells matches the expression pattern of β-CATENIN (Fig. 1C, C’), and CLAUDIN (Fig. 1D, D’) at E8.5, and E9.0 (Fig. 1F-G’). SOX9 was used as a marker of pre-EMT and migratory CNCCs in mouse embryos at E8.5 (Fig. 1B-D, red arrow). TWIST1 apical (arrowhead) is also observed at the dorsal edges of the neural tube just before fusion (Fig. 1H-H’). Notably, *Twist1* null embryos have complete neural tube closure defects (Fig. 1J, white arrow) compared to WT littermate (Fig. 1I, white arrow) at E11.5. In *Twist1* null embryos, the neural plate invaginated at the midline and lateral neural folds elevated towards the dorsal midline but the lateral folds failed to close properly. To determine whether a similar pattern of TWIST1 apical expression and adherens junction proteins might have a biological function (Fig. 1E’, F’), we performed co-immunoprecipitation (Co-IP) for TWIST1 to identify the protein interactors at E9.0-9.5. The Co-IP data shows that TWIST1 protein interacts with α/β/γ-CATENINS during neural tube development (Fig. 1K). Furthermore, β-CATENIN apical expression detected in WT was diffused in *Twist1^cko/−^* embryo and became mostly cytosolic (Fig. 1L-M’). To validate the IF staining data, whole mount *in situ* hybridization for *Twist1* mRNA shows weak expression at the dorsal edges of neural plate of E8.5 (Fig. 1N, N’; Fig. S2 and S3) and neural folds of E9.5 embryos (Fig. 1O, O’; Fig. S4). We also tested the quality and specificity of the anti-TWIST1 monoclonal antibodies in *Twist1* null embryos. Histological staining shows the neural tube closure defects in *Twist1* null compared to wild type embryos at E10.5 (Fig. S5A-B’). Notably, no signal for TWIST1 protein was detected in the apical side of neural tube or migratory mesenchymal cells of a *Twist1* null embryo in single and double staining with anti-IRF6 antibodies compared to a wild type embryo (Fig. S5C-F’).

**Fig. 1.**
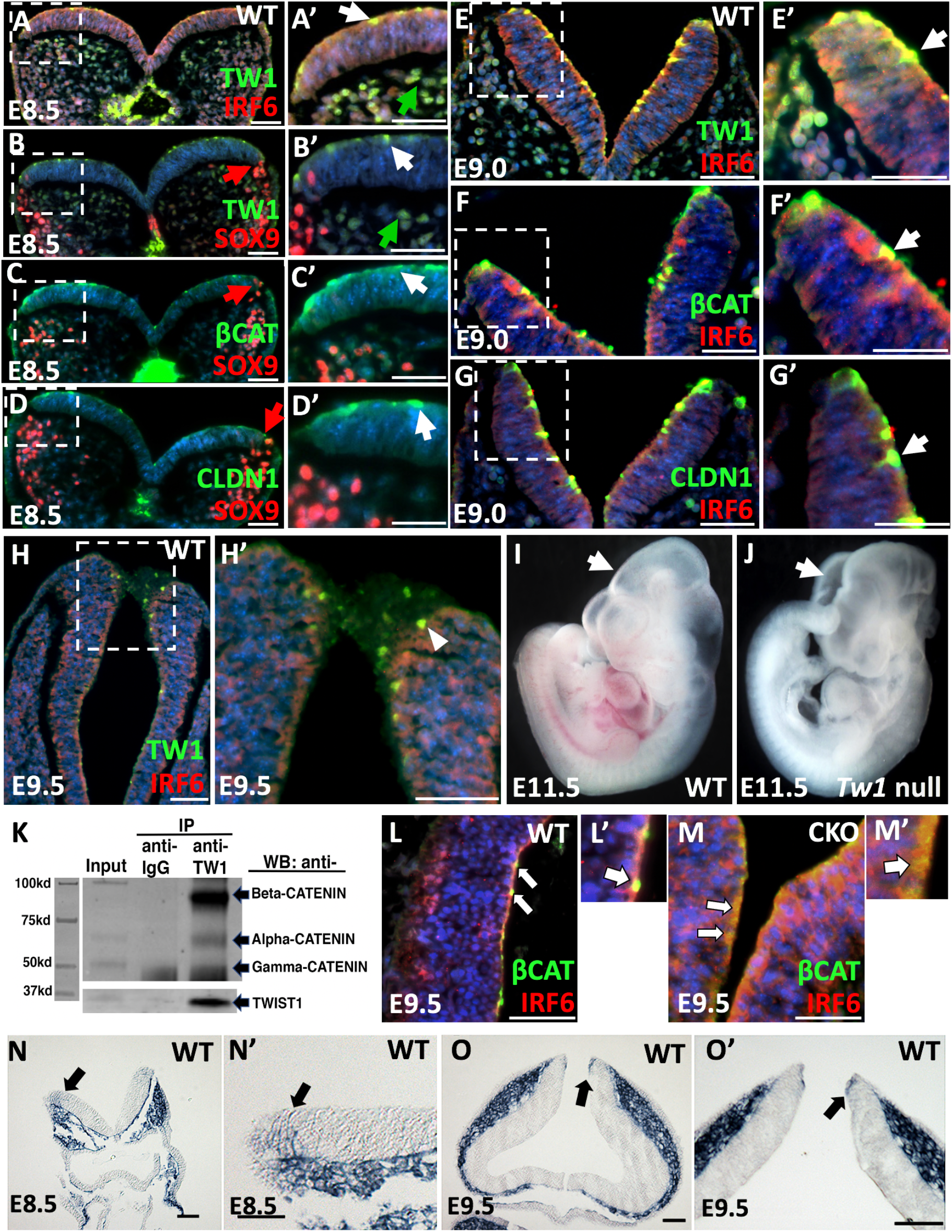
Spatiotemporal expression of TW1, IRF6, and other genes involved in neural tube development. (A, A’) IRF6 is present in the entire neural plate and TWIST1 is expressed in specific locations along the apical neural plate and in the adjacent mesenchymal cells at E8.5. (B, B’) SOX9 expression indicates NCCs and is seen in migrating cells as well as a few cells in the neural plate borders. (C, C’) β-CATENIN and (D, D’) CLAUDIN1 are expressed in specific locations along the apical side of neural plate similar to TWIST1. (E, E’). At E9.0 TWIST1 and IRF6 are expressed in similar locations as they were at E8.5. (F, F’) β-CATEININ and (G, G’) CLAUDIN1 are expressed in specific points along the apical side of neural folds at E9.0. (H) Expression of TWIST1 and IRF6 was also detected during closure of neural tube. TWIST1 is detected at the dorsal edges of neural tube (H, H’). (I) Wild-type embryos have closed neural tubes at E11.5, yet (J) *Twist1* null neural tubes are completely open. (K) The Co-IP assay shows enrichment of α/β/γ-CATENINS in the protein pulldown using TWIST1 monoclonal antibodies. Total proteins were extracted from hindbrain tissues protein of WT embryos. TWIST1 protein is detected in the input extract and is enriched in the pulldown extract using TWIST1 antibodies. (L, L’) β-CATENIN is detected apically in focal points of the dorsal cells of the neural tube and IRF6 is expressed throughout the neural tube at E9.5. (M, M’) β-CATENIN is detected in the cytosol of the dorsal cells of *Twist1^cko/−^* neural tube, while IRF6 expression is more prominent in the neural tube of *Twist1^cko/−^* embryo. (N-O’) *Twist1* mRNA is expressed in the dorsal neural folds of hindbrain at E8.5 and E9.5 by *in situ* hybridization. Scale bars represent 50 μm.

### Tracing CNCC migration in WT and *Twist1^cko/−^* using *ROSA26^Tm1^*; *Wnt1-Cre* and *Wnt1-Cre2* mouse line

To determine the role of *Twist1* in CNCCs, we used *Wnt1-Cre* and *Wnt1-Cre2* to conditionally knock out *Twist1* in pre-EMT CNCCs. For cell tracing, we used *R26^Tm1^* reporter gene and whole mount staining was performed in WT and *Twist1^cko/−^* using X-gal assay to detect CNCCs in blue. At E9.5, CNCCs were detected in the cephalic flexure of the midbrain, frontonasal process, and pharyngeal arches of WT embryos (Fig. 2A), and similar expression pattern was detected in WT at E11.5, except a stronger staining in the rhombic lips (Fig. 2B; Fig. S6A, A’). At E15.5 migratory CNCCs were observed in the otic pinna and frontal orofacial regions (Fig. 2C). In the mutant *Twist1^cko/−^* embryos (Fig. 2Aa, Bb, Cc; Fig. S6B, B’), weaker blue staining was observed in the frontonasal and pharyngeal processes, while stronger staining was detected in the mid- and hindbrain regions at E9.5, 11.5, and 15.5 suggesting a reduction in the number of migratory CNCCs towards frontonasal and pharyngeal processes. We noticed that the first pharyngeal arch of *Twist1^cko/−^* embryos was greatly reduced at E11.5, and the *Twist1^cko/−^* embryos displayed brain abnormalities at E15.5 (Fig. 2Bb, Cc). The recently generated *Wnt1-Cre2* mouse line was used to validate the CNCC-specific deletion by the *Wnt1-Cre* mouse line that has midbrain developmental abnormality as previously reported (Lewis et al., 2013). *Wnt1-Cre2* line showed a similar staining pattern of migratory CNCCs to *Wnt1-Cre* line (Fig. 2D, E, F), and the craniofacial phenotype associated with *Twist1* CKO in CNCCs (Fig. 2Dd, Ee, Ff). At E12.5 and E14.5, a dorsal image from the whole mount staining of WT embryos showed blue staining in the hindbrain and in the trunk neural tube (Fig. 2G, black arrow, Fig. 6A’). Similarly, blue staining was observed in *Twist1^cko/−^* embryos showing expanded staining in the mid- and hindbrain compared to the WT embryo (Fig. 2Gg, black arrow; Fig. S6B’). The embryos with compound alleles of *Twist1^cko/−^* and *Irf6^+/−^* presented severe hemorrhaging in the frontonasal process and craniofacial abnormalities compared to WT at E11.5 (Fig. 2H, Hh; Fig. S6C-E). Stereomicroscope images were taken from a WT embryo showing normal craniofacial development at E17.5 (Fig. 2I), and *Twist1^cko/−^* embryos showing severe exencephaly and abnormal frontonasal processes (Fig. 2Ii). Skeletal staining of *Twist1^cko/−^* embryos showed a loss of the cranial, frontonasal and maxillary bone as well as severe mandibular hypoplasia i (Fig. 2Jj) compared to WT littermates (Fig. 2J). To further determine the role of *Twist1* in the neural tube, hematoxylin and eosin (H&E) staining was performed in WT and *Twist1^cko/−^* embryos. H&E staining showed normal neural tube formation in WT (Fig. 2K, K’), while *Twist1^cko/−^* has multiple dorsolateral bends and expansion of the neural tube (Fig. 2L, L’). Histological staining of older WT embryos shows a normal development of the cephalic flexure of the midbrain and hindbrain neural tube at E10.5 (Fig. 2M, M’; Fig. S6A”-A””), while *Twist1^cko/−^* shows abnormal patterning of the cephalic flexure of midbrain and hindbrain neural tube (Fig. 2N, N’, black arrows; Fig. S6B”- B””). This data demonstrates that *Twist1* is crucial for the neural tube formation and migration of CNCCs towards the frontonasal processes and pharyngeal arches.

**Fig. 2.**
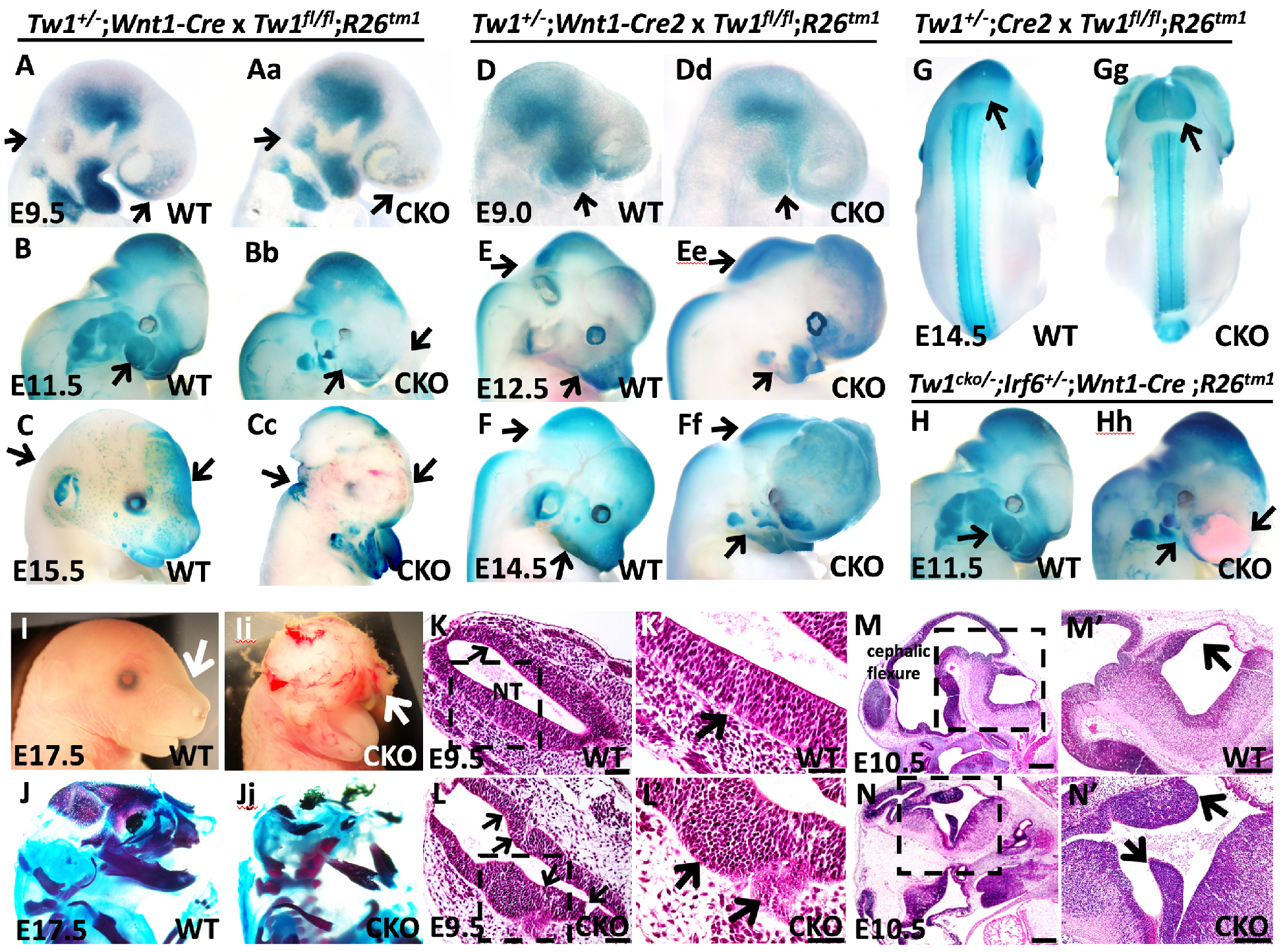
CNCC migration in WT and *Twist1^cko/−^* using an *in vivo* tracing system. (A) At E9.5, CNCCs stained in blue are detected in cephalic flexure, frontonasal process, and pharyngeal arches in WT. (Aa) Less staining of migratory CNCCs is observed in frontonasal and pharyngeal arches of *Twist1^cko/−^* embryos, while stronger blue staining is detected in the hindbrain regions. (B, Bb) The mandibles of *Twist1^cko/−^* embryos are smaller (agnathia) than WT, and the blue staining of CNCCs is weaker in the frontal craniofacial processes. (C, Cc) At E15.5, the *Twist1^cko/−^* embryo lacks proper craniofacial development and displayed exencephaly, and retained blue staining at the hindbrain. (D-Gg) *R26^Tm1^; Wnt1-cre2* cell tracing from E9.5 to E14.5 in WT and *Twist1*^cko/−^ embryos show a similar blue staining pattern to *Wnt1-Cre*. (H, Hh) *Irf6^+/−^; Tw1^cko/−^; Wnt1-cre* embryo shows under-developed mandible, severe forebrain hemorrhage, and darker staining at the hindbrain regions at E11.5. (I, Ii) Macroscopic images of WT and *Twist1*^cko/−^ embryos at E17.5. (Ii) Mutant embryo shows severe exencephaly and loss of craniofacial structures. (J-Jj) Skeletal staining of WT compared to *Twist1^cko/−^* showing loss of most craniofacial bone (red) and nasal cartilage (blue). (K, K’) Hematoxylin and eosin staining shows normal neural tube formation in WT. (L, L’) *Twist1^cko/−^* neural tube has multiple bends and lateral expansion. (M, M’) Histological staining shows normal development of cephalic flexure of midbrain and hindbrain neural tube in WT embryos. (N, N’) The abnormal pattern of midbrain and hindbrain is detected in *Twist1^cko/−^* with unorganized protrusions in hindbrain (N, N’, black arrows). Scale bars represent 100 μm.

### CNCC delamination and migration in WT and *Twist1^cko/−^* neural tube explants

The reduction in CNCC population in the frontonasal processes and pharyngeal arches of *Twist1^cko/−^*; *R26^tm1^* embryos leads to prenatal lethality due to severe exencephaly and lack of craniofacial bone and cartilage (Fig. 2Cc, Ff, Ii). We further investigated why a loss of *Twist1* pre-EMT neuroectodermal cells leads to a severely abnormal brain and craniofacial structures, even though a decent amount of CNCCs migrated towards the frontonasal and pharyngeal processes. We also asked why migratory CNCCs did not form craniofacial bone and cartilage (Fig. 2Jj). To answer these questions, we developed a neural tube explant system using a dual reporter transgenic mouse line (*ROSA26^tm4.ACTB-tdTomato-EGFP^*), which allows us to capture cell morphogenesis and migration of somatic and CNCCs. The illustrative diagram depicts the procedure of how the neural folds were dissected from embryos at E8.5 and the neural fold of the mid- and hindbrain was cultured overnight on a culture plate coated with collagen type I (Fig. 3A). We used this system to uncover the function of *Twist1* in pre-EMT CNCCs, wherein lack of its expression in the neuroectodermal cells and pre-EMT neural crest can explain the phenotype observed in *Twist1* null and CKO in CNCCs. The cells expressing the *Wnt1-Cre* transgene excise the membrane-associated *mTomato* reporter gene, allowing instead for the expression of *GFP* reporter gene. In WT explants, individual migratory CNCCs moved away from the dorsolateral sides of the neural folds (Fig. 3B, B’; Time-lapse movie 1) and the migratory mesenchymal cells formed long protrusions (inset in Fig. 3B’; Time-lapse movie 2). In the *Twist1^cko/−^* explant, a considerable amount of *Twist1*-deficient neuroectodermal cells remained within the neural folds and expanded ventrally (Fig. 3C; Time-lapse movie 3), and the majority of the detached CNCCs did not transition into individual mesenchymal cells but instead maintained their cell-cell adhesion and epithelial morphology (Fig. 3C’, inset in C’; Time-lapse movie 4). Quantitative measurements of the total number of CNCCs that migrated away from the neural tube were analyzed for three biological replicates of each genotype. We detected approximately a 50% reduction in the total number of cells in mutant explants (Fig. 3D), and the average distance of all migratory CNCCs was reduced around 68% in mutant explants (Fig. 3E). We also measured the speed mean of individual CNCCs in WT and *Twist1^cko/−^*. To determine the mean speed of individual CNCCs, we tracked the CNCCs as they moved away from the neural tube explants of WT and *Twist1^cko/−^* (Fig. 3F, F’, G, G’). Micrographs show individual CNCC migration paths during time-lapse imaging (Fig. 3F’, G’, white lines). Wild type CNCCs mean speed increased by 50% in comparison to *Twist1^cko/−^* (Fig. 3H).

**Fig. 3.**
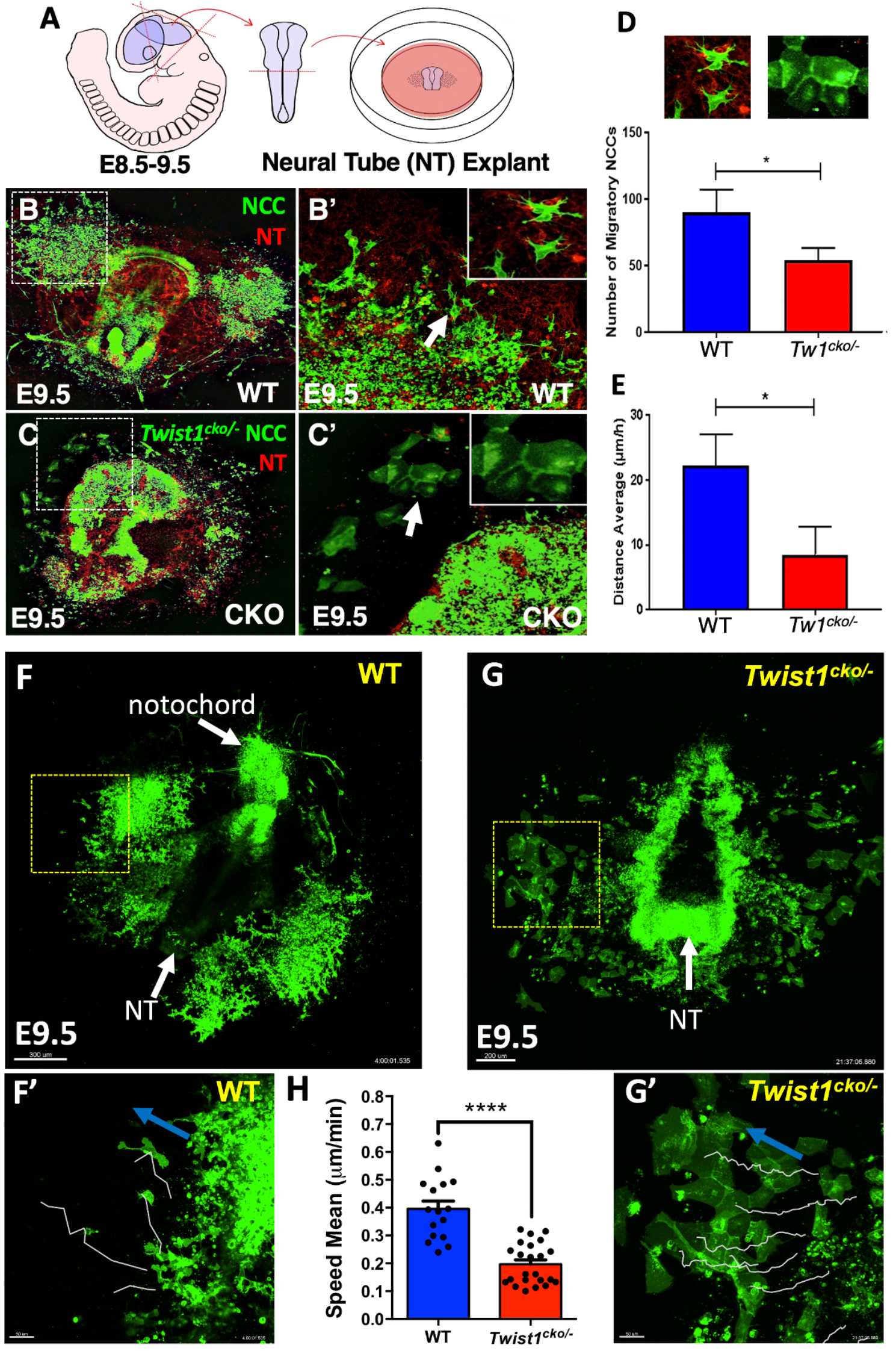
Neural tube (NT) explant culturing for time-lapse imaging of WT and *Twist1^cko/−^* CNCCs during cell formation and migration. (A) An illustrative diagram showing the NT explant (E8.5-9.5) preparation process. (B-C’) Confocal time-lapse images of NT explants at E9.0. Green shows migratory CNCCs in WT (B, B’), and *Twist1^cko/−^* neural tube in which less CNCCs delaminated from the tube and migratory cells maintain their cell-cell adhesion (C, C’). Many pre-EMT CNCCs remained in *Twist1^cko/−^* neural tube and expanded ventrally (C, C’), white box and arrow indicate the magnified area location. (D) Quantification of total CNCCs that migrated away from the neural tube, and Individual cell migration paths (white lines) are shown for WT and *Twist1^cko/−^* explants. (E, F) Show the entire mouse embryo explant for WT and Twist1^cko/−^, and (E’,F’) show the magnified areas specified by a box in E and F. Blue arrows show the direction of cell migration for each explant. (H) Average speed mean migration (µm/min) of WT and *Twist1^cko/−^* CNCCs. (*) Indicates a statistically significant difference (*p* < 0.05). Scale bars represent 300 μm for F, 200 μm for G, and 50 μm for F’ and G’.

### Neural tube phenotype in *Irf6* null embryos

IRF6 is expressed in the non-neural and neural ectoderm during neural tube formation. Its expression is relatively stronger in neural ectoderm than non-neural ectoderm (Fig. 1E, Fig. S1E, F, G). We hypothesized that a complete loss of *Irf6* affects the structural integrity of the neural tube, leading to craniofacial abnormalities in *Irf6* null embryos. At E10.5, we noticed that the cellular structures of the dorsal edges of neural tube and apical membrane were disorganized in *Irf6* null compared to WT littermates (Fig. 4A-B”). Abnormal expansions from the brain cortex also were detected in *Irf6* null embryos compared to WT littermates (Fig. 4C-D’). We checked the expression of SOX9, which is expressed in the neuroepithelial progenitors of glial cells (Farrell et al., 2011; Vogel and Wegner, 2021), and F-ACTIN in WT and *Irf6* null embryos. SOX9 positive cells were detected in the dorsal half of the neural tube (Fig. 4E, E’, white arrow). However, we found that the organization of SOX9 positive cells was altered and large intercellular gaps among SOX9 positive cells were detected in *Irf6* null neural tube (Fig. 4F, F’, white arrows). A continuous expression of F-ACTIN was detected in the apical cells of neural tube dorsal edges of WT (Fig. 4E’), while the apical membrane at the neural tube edges was discontinuous in *Irf6* null embryos (Fig. 4F”). We also investigated the expression of non-neural ectoderm marker, AP2α, to uncover whether a loss of *Irf6* alters the expression of AP2α at the neural tube junction with non-neural ectoderm. We noticed that the number of AP2α positive cells at the neural tube dorsal edges increased in *Irf6* null embryos compared to WT littermates in four different biological replicates of each genotype (Fig. 4G, H). We quantified the number of AP2α positive cells in the dorsal edges of neural tube and observed a significant increase in *Irf6* null compared to WT (Fig. 4I).

**Fig. 4.**
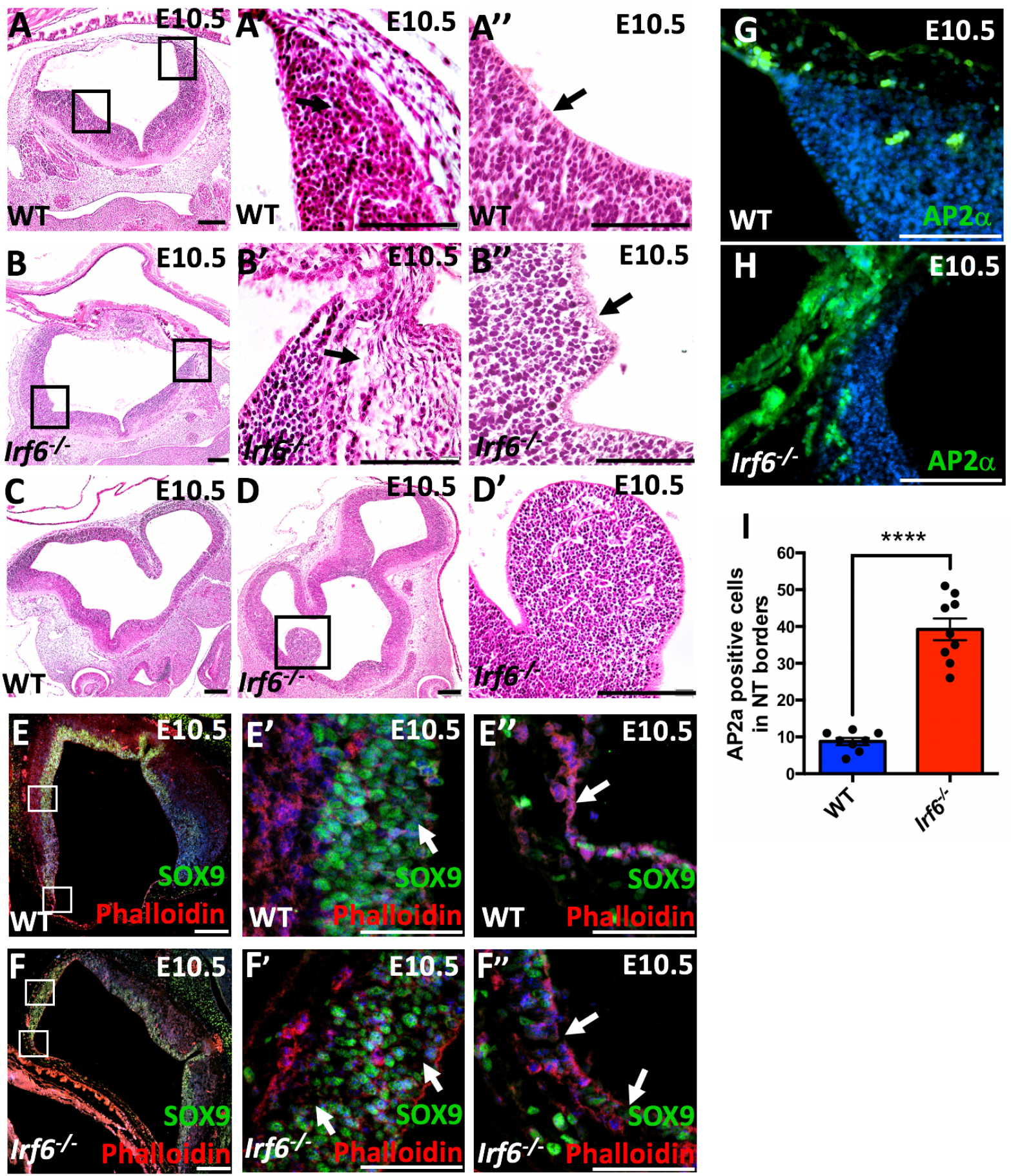
Neural tube phenotype in *Irf6* null embryos. (A-A”) Histological staining of WT embryonic sections shows the morphology of the neural tube (A), the neural tube dorsal edges (A’), and neural tube apical cells (A”) at E10.5. (B-B”) Histological staining of *Irf6* null embryonic sections shows the neural tube morphology (B), neural tube edge with less organized neuroepithelial cells and larger intercellular space (B’), and neural tube epical layers (B”). The apical membrane is expanded and shows a wavey pattern (B”). (C) Histological staining of embryonic section of WT embryos shows the forebrain morphology. (D-D’) Abnormal protrusions from the brain cortex are detected in *Irf6* null embryos compared to WT littermates. (E-F”) Expression of SOX9 (gliogenesis marker) and F-actin was detected in WT (E-E”) and *Irf6* null embryos (F-F”). The organization of SOX9 positive neuroepithelial cells is altered with increased intercellular space between the cells in *Irf6* null embryos (F, F’) compared to WT (E, E’). The integrity of the apical membrane at the neural tube is altered in *Irf6* null embryonic section (F”) compared to WT embryo (E”). (G) Immunostaining of AP2α in green shows expression in ectoderm and the junction between non-neural and neural tube edge in WT. (H) Increased expression AP2α is detected in the neural tube edges and the junction between non-neural ectoderm and neural tube edge. (I) The quantification of the AP2α positive cells at the neural tube edges shows a significant increase in *Irf6* null embryos compared to WT littermates. Scale bars represent 100 μm.

### Expression pattern and level of epithelial genes and WNT3 in neural tube and migratory mesenchymal cells

Based on the CNCC-derived tissue phenotypes observed in *Twist1^cko/−^*, we investigated the expression pattern and level of epithelial genes and WNT3 involved in cell differentiation and adhesion to determine why a loss of *Twist1* in delaminated CNCCs maintained their epithelial morphology. Using IF staining, we first looked *in vivo* at the expression of IRF6 and TWIST1 in WT and *Twist1^cko/−^* in more than four biological replicates for each gene. In WT embryos, IRF6 was mainly detected in the neural tube while TWIST1 was highly expressed in migratory mesenchymal cells that were detached from the neural tube and migrated towards frontonasal and pharyngeal prominences at E9.5-E10.5 (Fig. 5A, A’; Fig. S7A-C). Ectopic IRF6 expression in partially detached mesenchymal cells in *Twist1^cko/−^* embryo was detected in a cluster of cells along the neural tube (Fig. 5B-B’, white arrow). No expression of E-CADHERIN (E-CAD) was present in mesenchymal cells along the neural tube in WT embryos (Fig. 5C, C’). However, staining for E-CAD in *Twist1^cko/−^* embryos showed that the partially attached mesenchymal cells along the neural tube express ectopic E-CAD (Fig. 5D, D’). WNT3 expression was detected at the junction of neural tube dorsal edges and in a few mesenchymal cells in WT embryo (Fig. 5E, E’), while decreased and expanded expression of WNT3 in the neural tube was detected in *Twist1^cko/−^* (Fig. 5F-F’). We also looked at the expression of tight junction proteins, OCCLUDIN and ZO1, *in vivo* and in neural tube explants. OCCLUDIN is highly expressed at the junction of neural tube dorsal edges of WT embryos (Fig. 5G, G’), while almost no expression of OCCLUDIN was detected at the unfused junction of the neural tube dorsal edges of *Twist1^cko/−^* embryos (Fig. 5H, H ‘). In neural tube explants, an interspaced and broken signal of ZO1 was seen between the migratory mesenchymal cells that migrated away from the neural tube explant in WT (Fig. 5I, I’), while a continuous junctional signal of ZO1 was observed among detached mesenchymal cells from the neural tube explant of *Twist1^cko/−^* (Fig. 5J, J’). In addition, we looked at the expression of a mesenchymal marker, VIMENTIN, *in vivo* and in neural tube explants and observed a disrupted expression pattern in the mesenchymal cells in *Twist1^cko/−^* embryos and neural tube explant compared to WT (Fig. S7D-G’). mRNA expression of *Twist1* and *Irf6* in the hindbrain and first pharyngeal arch was also measured in 3 pooled biological and 4 technical replicates of WT and *Twist1^cko/−^*embryos at E9.5. As expected, *Twist1* expression was significantly reduced in *Twist1^cko/−^* embryos, and a slight increase was seen in the *Irf6* mRNA of *Twist1^cko/−^* (Fig. 5K). mRNA encoding *E-Cad*, *N-Cad*, and *β-Catenin* were also measured in WT and *Twist1^cko/−^* (Fig. 5L). *E-Cad* expression was significantly increased in *Twist1^cko/−^* (Fig. 5L), which is consistent with the ectopic expression of E-CAD in partially detached mesenchymal cells (Fig. 5D, D’).

**Fig. 5.**
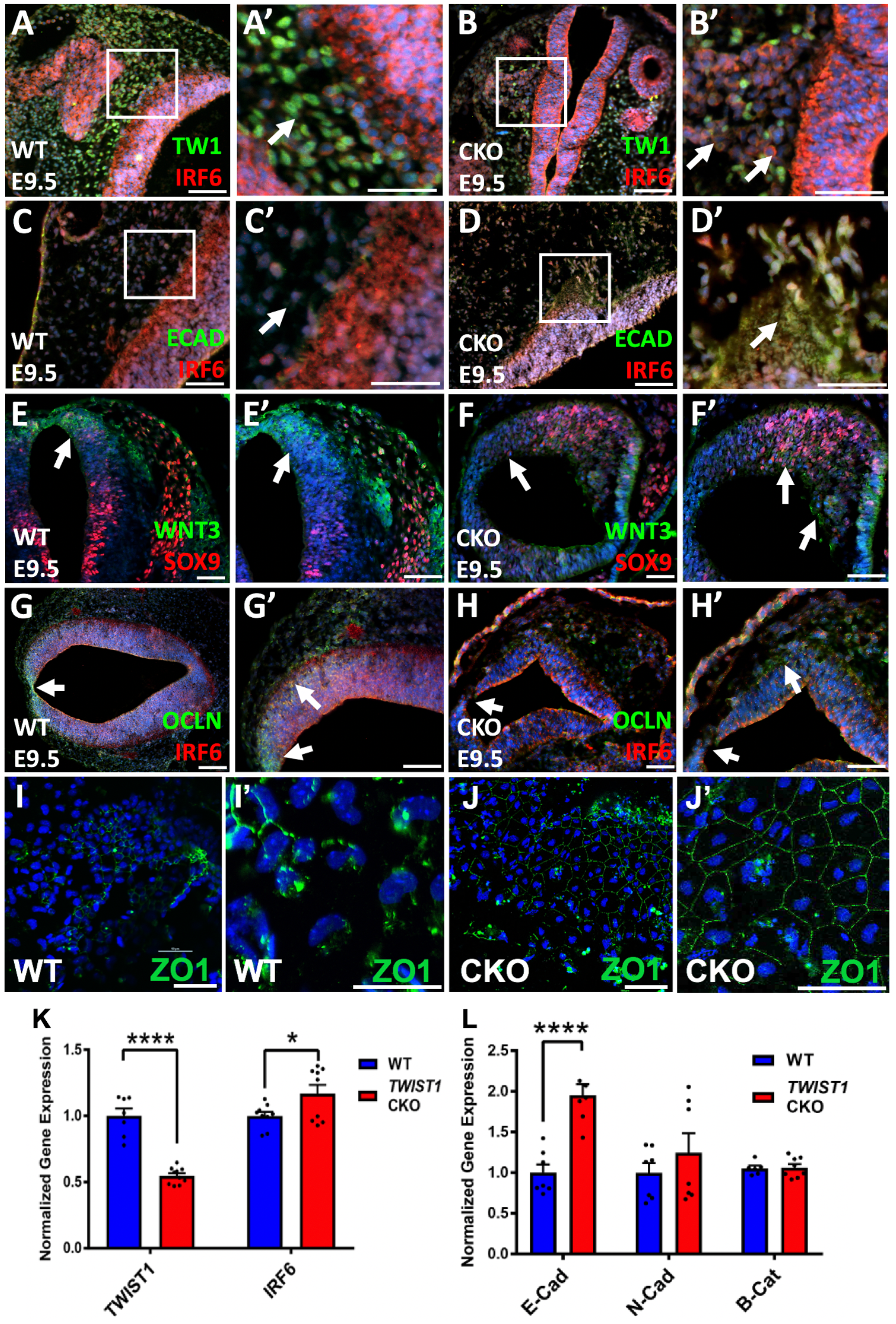
Expression of IRF6, TWIST1 and other factors during neural tube and CNCC formation. (A, A’) IRF6 is detected in the neural tube while TWIST1 is expressed in migratory CNCCs at E9.5. (B, B’) IRF6 expression in *Twist1^cko/−^* embryos is detected in the partially delaminated CNCCs along the neural tube. (C, C’) Staining for E-CADHERIN in wild type shows expression in ectoderm and apical regions of the neural tube, but at a low level. (D, D’) In *Twist1^cko/−^* embryos, ectopic E-CADHERIN expression is detected in the partially detached and migratory mesenchymal cells. (E, E’) WNT3 is observed at the dorsal side of the neural tube in wild type embryos. (F, F’) WNT3 expression is decreased at the dorsal side of neural tube and a signal is detected ventrally in *Twist1^cko/−^* embryos. (G, G’) OCCLUDIN is highly expressed at the junction of the neural tube edges. (H, H’) OCCLUDIN signal is reduced at the junction of the neural tube edges of *Twist1^cko/−^*. (I, I’) ZO1 is detected as an interspaced and broken signal in WT CNCCs. (J, J’) ZO1 is expressed as a junctional signal between the detached CNCCs of *Twist1^cko/−^* explants. (K, L) Expression of *Twist1* and *Irf6* mRNA in the hindbrain and lower jaw was measured in WT and *Twist1^cko/−^* embryos. (K) *Irf6* mRNA expression shows a slight increase in the *Twist1^cko/−^*. (L) mRNA of *E-Cad*, *N-Cad* and *β-Cat* were measured in WT and *Twist1^cko/−^*. *E-Cad* expression is significantly increased in *Twist1^cko/−^*. The *E-Cad* mRNA level is increased similar to the ectopic expression of E-CAD in detached and migratory CNCCs (D). Scale bars represent 50 μm.

### Quantitative levels of kinases, signaling molecules, and epithelial and mesenchymal marker genes

The expression of genes encoding kinases (*Akt1, Akt2, Pi3k, Erk1, Ck2, cSrc*), epithelial markers (*E-Cad, N-Cad, Occludin*), neural and non-neural ectoderm markers and CNCC specification drivers (*Snail2, Msx1, AP2α, Hand2, Sox2, Sox10, HoxD10*), signaling molecules (*Yap, Wnt1, Wnt3, Jag1, mTor*), and actomyosin remodeling regulators (*β-Catenin, RhoA, RhoC, Arhgap29*) was measured in dissected tissues of the hindbrain and first pharyngeal arch (Fig. 6; Fig. S8). These genes were selected based on previous developmental and cancer studies on TWIST1 and its associated-regulatory pathways. By comparing the expression of potential target genes in WT versus *Twist1^cko/−^* embryos, we anticipated to uncover the differentially expressed genes involved in *Twist1*-dependent cellular function. The mRNA expression of *Akt2*, *Snail2*, *Msx1*, *Hand2*, *Sox10*, *Yap*, *β-Catenin, RhoA* and *Arhgap29* was significantly reduced in *Twist1^cko/−^* embryos at E10.5 (Fig. 6A, E, C, G, I, and Fig. S8A). Protein analysis by western blot also showed decreased levels of AKT1/2/3, SNAIL2, and WNT3 in *Twist1^cko/−^* embryos at E10.5 (Fig. 6B, F, H). Although the *in vivo* ectopic expression of E-CAD was observed in partially delaminated mesenchymal cells at E9.5, and the mRNA level was significantly increased in *Twist1^cko/−^* at E10.5, the total protein level in the hindbrain and first pharyngeal arch was not different between CKO and WT due to eventually the contribution of other tissues such as ectodermal and endodermal cells (Fig. 6D).

**Fig. 6.**
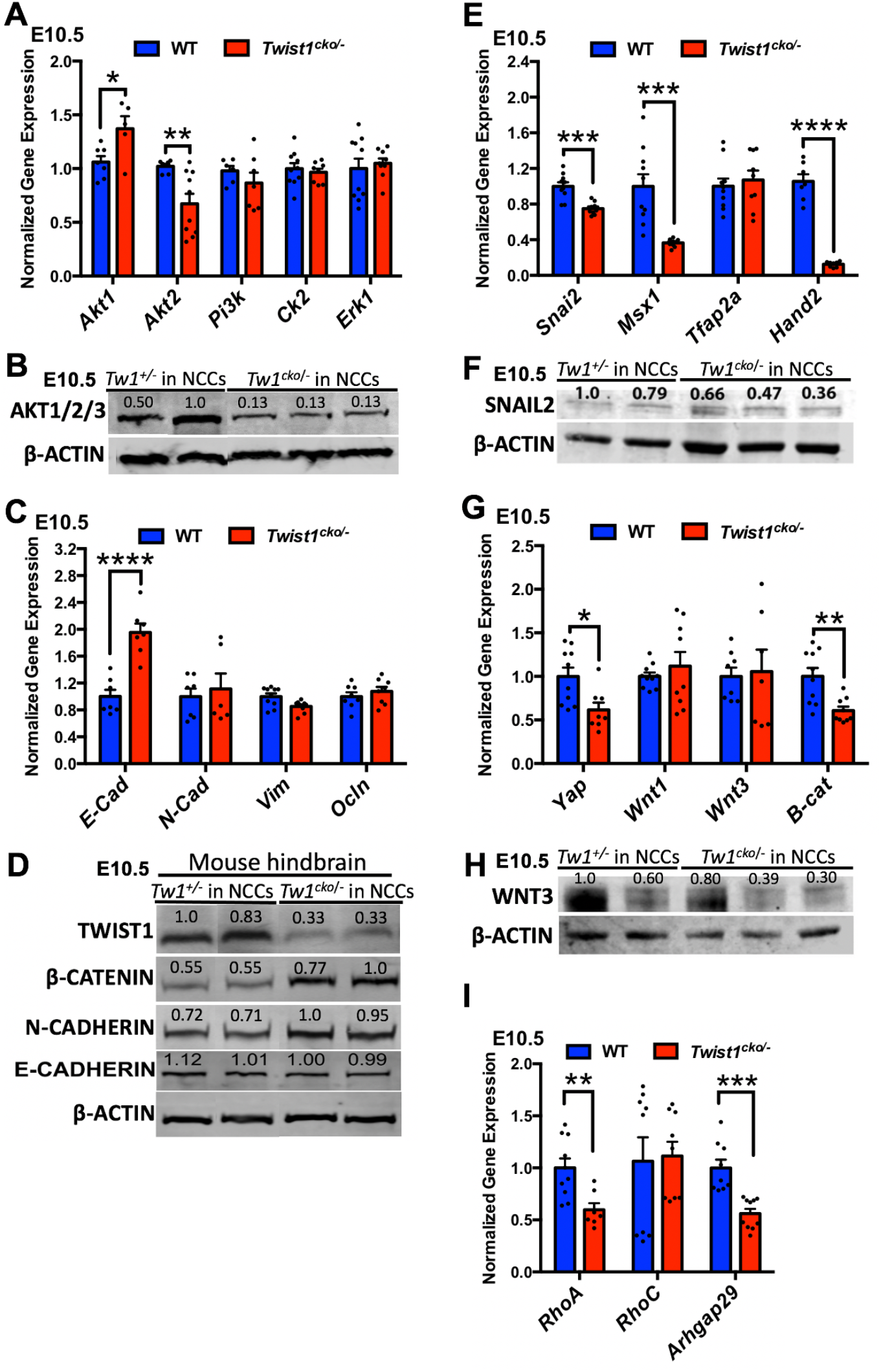
Differentially expressed genes in *Twist1^cko/−^ in-vivo*. (A) Comparison of the expression of *Akt1*, *Akt2,* and *Pi3k, Ck2* and *Erk1* mRNA between WT and *Twist1^cko/−^*. (B) Protein analysis by western showed a reduction in the kinase expression in mutant embryos at E10.5. (C) Comparison of the expression of *E-Cadherin* (adherens junction marker), *N-Cadherin*, *Vimentin* (mesenchymal marker), and *Occludin* mRNA in WT and *Twist1^cko/−^*. (D) Protein analysis by western showed that adherens and tight junction proteins are increased in mutant embryos atE13.5. (E) Comparison of the expression of *Snai2, Msx1, Tfap2a and Hand2* mRNA between WT and *Twist1^cko/−^*. (F) Protein analysis by western showed a reduction in the expression of SNAI2 transcription factor in mutant embryos at E10.5. (G) Comparison of the expression of *Yap*, *Wnt1*, *Wnt2*, *Wnt3* and *β-Catenin* mRNA in WT and *Twist1^cko/−^*. (H) Protein analysis by western showed that WNT3 protein is decreased in mutant embryos at E13.5. (I) Comparison of the expression of *RhoA, RhoC, and Arhgap29* mRNA in WT and *Twist1^cko/−^*. The level of *RhoA* and *Arhgap29* is significantly reduced in *Twist1^cko/−^* embryos compared to WT.

### TWIST1 nuclear expression and phosphorylation *in vivo* and in O9-1 cell line

Previous studies demonstrated that 6 out of 9 predicted residues of TWIST1 are phosphorylated in human cancer cell lines. We mapped the conservation of the 6 phosphorylated serine and threonine residues in *Twist1* based on the bioinformatic analysis of vertebrate species and *D. melanogaster* (Fig. 7A). TWIST1 has 6 phospho-residues that are highly conserved in vertebrates, particularly T121 and S123 that are conserved all the way to *D. melanogaster*. The *in vitro* function of each phosphoresidue was based on previous cancer studies (Fig. 7A, bottom row) (Xue and Hemmings, 2012). Our data shows that some subpopulation in the dorsal edges of neural folds and in the pre-EMT CNCCs express TWIST1 mostly in the cytosol as per IF staining data (Fig. 7B-C’, arrows; Fig. S1A’, D, D’), and it becomes mainly nuclear in detached and migratory CNCCs at E9.5 and E10.5 (Fig. 5A, A’; Fig. 7D, D’, arrow). TWIST1 is predominantly expressed in migratory CNCCs during the development of frontonasal and pharyngeal arch processes (Fig. S7A, B, C. Our *in vivo* data shows that TWIST1 protein is phosphorylated in mesenchymal cells at S42, S68, and S123 residues at E10.5. The phosphorylated form was detected after enrichment of TWIST1 by immunoprecipitation using monoclonal antibodies for TWIST1 and blotting against phosphorylated forms of TWIST1 using monoclonal anti-P-S68 and polyclonal antibodies counter to P-S42 and P-S123 (Fig. 7E). However, the expression of phosphorylated TWIST1 at S68 was weaker compared to S42 and S123. TWIST1 unphosphorylated and phosphorylated forms were also detected in the O9-1 cell line without enrichment of TWIST1 protein and using the monoclonal antibodies that can recognize both forms (Fig. 7F). To validate the TWIST1 phosphorylated band, the total protein was treated with protein phosphatase 2 (PP2) to remove the phosphate group from phosphorylated proteins. The TWIST1 phosphorylated band was decreased without affecting the band intensity of the non-phosphorylated band (Fig. 7F).

**Fig. 7.**
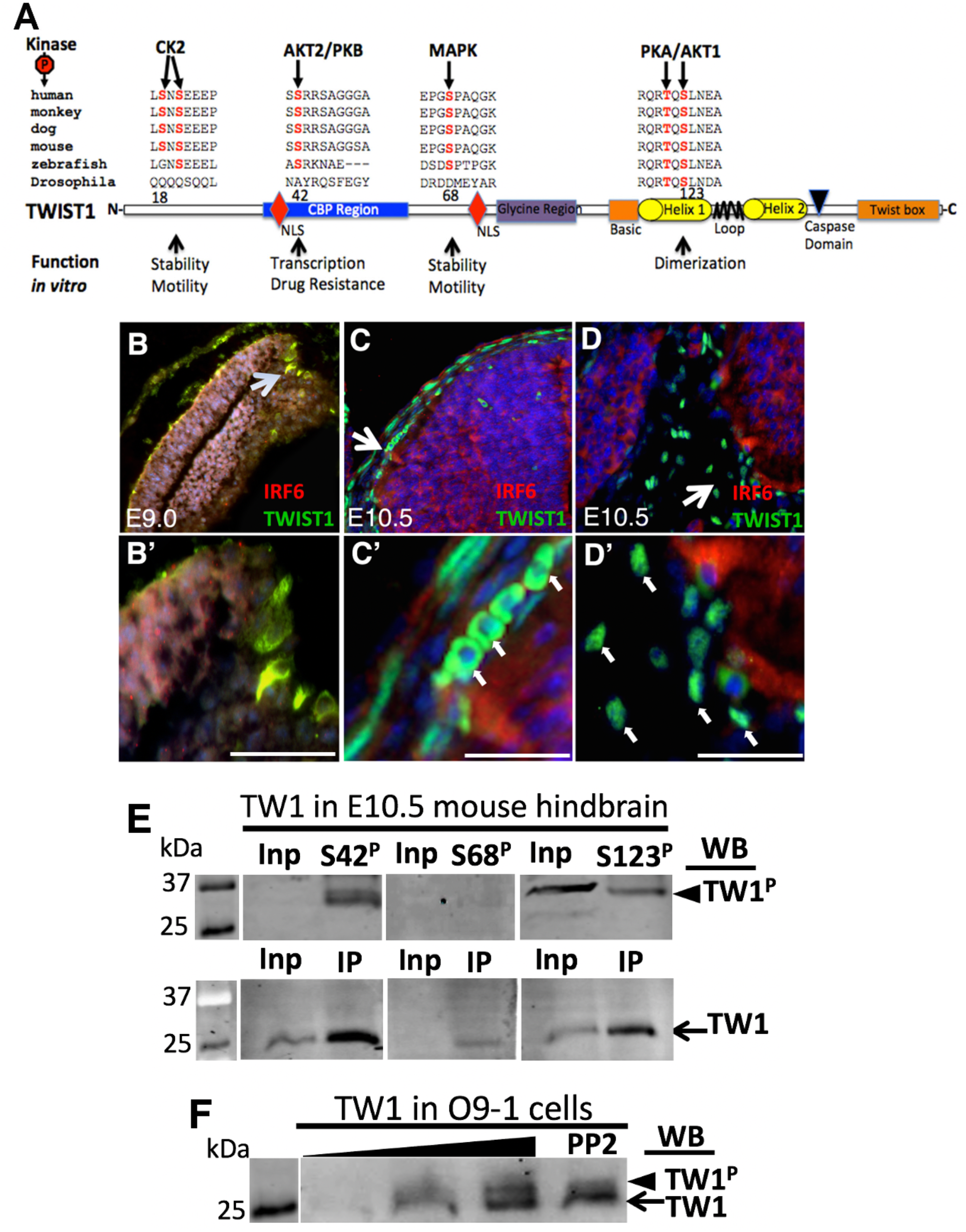
TWIST1 nuclear translocation and phosphorylation in CNCC. (A) Mapping of phosphorylation sites within *TWIST1* functional domains and their respective kinases in cancer cell lines. TWIST1 has 6 phospho-residues based on *in vitro* cancer studies. All are highly conserved, particularly T121 and S123. (B-D’) TWIST1 and IRF6 expression pattern in neural tube and CNCCs. (B-B’) At E9.0, TWIST1 is expressed in the cytosol of dorsal lateral cells of the invaginated neural plate. (C-C’) At E10.5, TWIST1 is mostly cytosolic in pre-delamination CNCCs. (D-D’) TWIST1 becomes nuclear when the CNCCs detach and migrate toward the pharyngeal arches. (E) Immunoblot showing TWIST1 phosphorylated bands of S42, S68 and S123 (arrowhead) using polyclonal rabbit antibodies before immunoprecipitation (Input) and after enrichment (IP) using monoclonal TWIST1 antibodies (bottom blot, row) in WT mouse embryo at E10.5. (F) TWIST1 unphosphorylated (bottom arrow) and phosphorylated forms (arrowhead) are also highly expressed in O9-1 cells. Scale bars represent 50 μm.

### Importance of TWIST1 phosphorylation in craniofacial tissues

To determine the role of TWIST1 posttranslational modifications in the neural tube and CNCC-derived tissues, we generated four phospho-incompetent mouse lines, including *Twist1^S18;20A^*, *Twist1^S42A^*, *Twist1^S68A^*, and *Twist1^T121A; S123A^* using CRISPR/CAS9 technology. The founders of *Twist1^S42A^* mice did not give any progeny, and all ten *Twist1^T121A; S123A^* founders died before reaching two months of maturity. We confirmed the genomic changes in 3 founders (2 males and 1 female) of *Twist1^S18I;20A/+^*, and four founders of *Twist1^S68A/+^* (2 males and 2 females). We backcrossed all phospho-mutant founders to C57BL/6J WT mice for 6 generations to avoid non-specific genomic changes or alterations. The Sanger sequence of F6 generations confirmed the changes of serine 18 (S18) to isoleucine and serine 20 (S20) to alanine, and in the second mouse line, serine 68 (S68) to alanine (Fig. 8A, B).

**Fig. 8.**
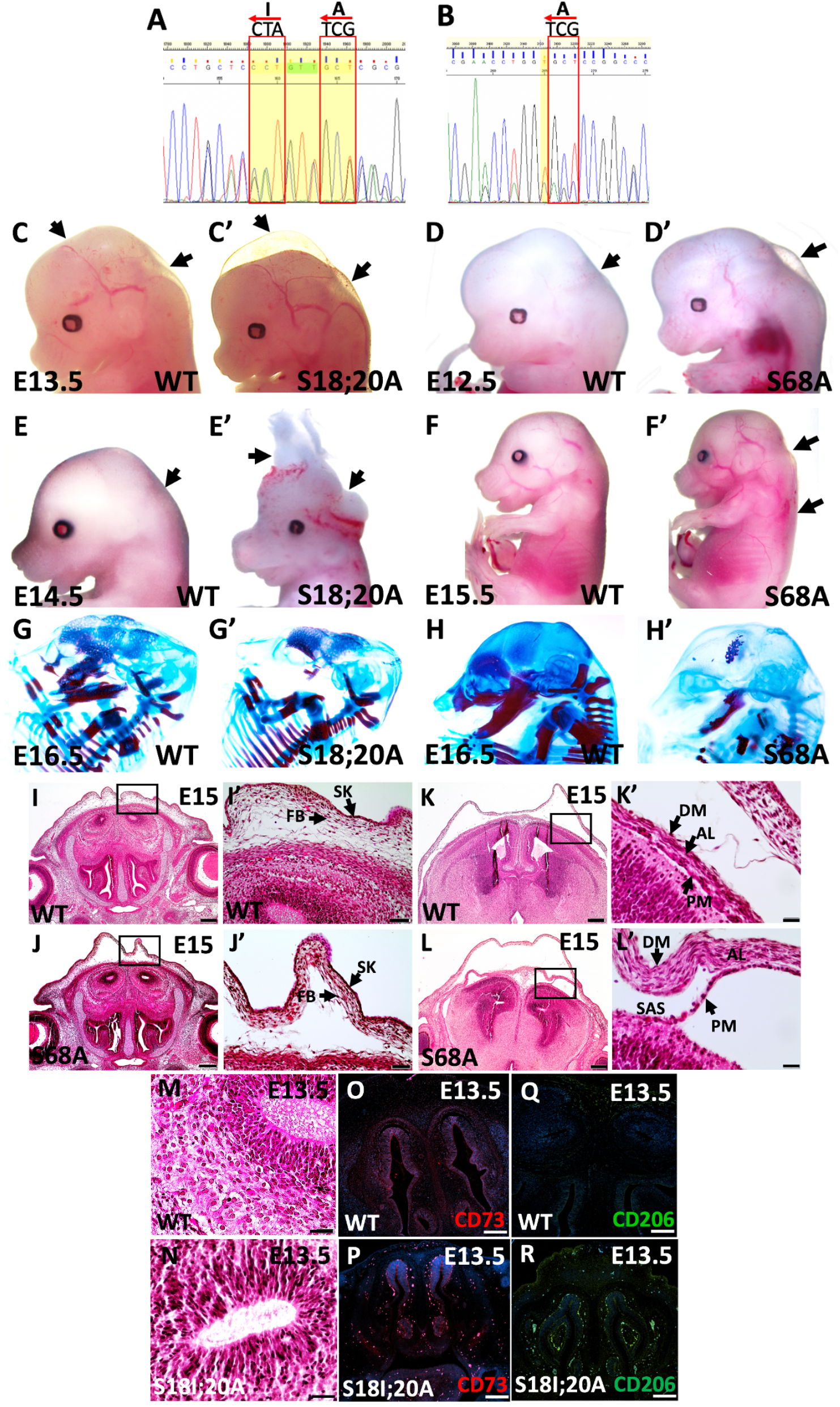
The *Twist1^S18;20A/S18;20A^* and *Twist1^S68A/S68A^* morphological and skeletal phenotype. (A)The DNA sequences show the changes in the serine residue S18/20 to isoleucine and alanine, and S68 to alanine. (C-F’) Both *Twist1^S18;20A/S18;20A^* and *Twist1^S68A/S68A^* mice show epidermal blebbing, severe edema along the neural tube, and neural tube defects. Skeletal staining of *Twist1^S18;20A/S18;20A^* and *Twist1^S68A/S68A^* mutant embryos shows a loss of craniofacial bone and a reduction in skull mineralization. (G, G’) *Twist1^S18;20A/S18;20A^* mutant embryo has no mandibular and maxillary bone, or the bone formation is delayed compared to WT. (H, H’) Skeletal staining of *Twist1^S68A/S68A^* shows a remarkable reduction in skull mineralization of the frontonasal and maxillary bone of most cranial and limb bone and cartilage at E16.5. (I-K’) Histological staining shows normal morphology of frontonasal structures of a coronal section of wild type embryo at E15. (J-L’). In *Twist1^S68 A/S68A^* embryo, histological staining shows subepidermal blebbing, detachment of fibroblast layers from brain and meninges abnormalities. (M, N) Histological staining in the brain of WT and *Twist1^S18;20A/S18;20A^* shows infiltration of lymphocytes in *Twist1^S18;20A/S18;20A^* compared to WT. (O, Q, P, R) IF staining shows no signal for CD73 and CD206 in WT, however, a strong staining was observed in the brain and around nasal cavities of *Twist1^S18;20A/S18;20A^*. Scale bars represent 100 μm.

The phenotypic characterization showed that the two phosphorylation sites, S18/20 and S68, are critical for TWIST1 activity in craniofacial tissue development. The two phospho-incompetent mouse lines have epidermal blebbing, severe edema along the neural tube, and significant neural tube defects in *Twist1^S18I;20A/S18I;20A^* compared to WT littermates (Fig. 8C-F’). The skeletal staining exhibited a remarkable loss of maxillary and mandibular bones in *Twist1^S18I;20A/S18I;20A^* and a reduction in skull mineralization of the frontonasal and maxillary bones in *Twist1^S68A/S68A^*, respectively (Fig. 8G’, H’), compared to WT controls (Fig. 8G, H). The histological staining of *Twist1^S68A/S68A^* head structures at E15 showed subepidermal blebbing, separation of fibroblast layers from the meninges, and a detachment of the meninges from the surface of the brain cortex (Fig. 8J, J’, L, L), compared to WT littermates (Fig. 8I, I’, K, K’). We also noticed lymphocyte infiltration in *Twist1^S18I;20A/S18I;20A^* brain tissues and around the frontonasal processes compared to WT embryos (Fig. 8M, N). To determine the types of the infiltrated cells, IF staining for the lymphocyte marker CD73 and B cell marker CD206 were utilized on tissues sections at E13.5. No signal was detected in the frontonasal sections of WT (Fig. 8O, Q), however, enrichment of both markers was noticed in the frontonasal tissues of *Twist1^S18I;20A/S18I;20A^* embryos (Fig. 8P, R).

Large numbers of embryos were collected from the two *Twist1* phospho-incompetent mouse lines to determine the ratio of genotype to phenotype and disease penetrance. The phenotype penetrance of the homozygous *Twist1^S18I;20A/S18I;20A^* and *Twist1^S68A/S68A^* mice was incomplete as shown in Fig. S9, because the percentage of homozygous embryos and pups was 23.5% and 21.4%, respectively, as shown in Fig. S9. However, the homozygous *Twist1^S18I;20A/S18I;20A^* and *Twist1^S68A/S68A^* embryos with phenotypes constitutes 11.5% and 14.6% of the total percentage, respectively, suggesting an incomplete penetrance of the phenotype (Fig. S9).

### Interaction between *Twist1* and *Specc1l* in CNCCs

Many of the tested cytoskeletal remodeling regulators were significantly reduced at the mRNA level in *Twist1^cko/−^* embryos, while the *in vivo* expression of adherens junction proteins by IF is altered. We focused on *Specc1l* because of the phenotypic overlap in craniofacial disorders in humans and mice and the importance of this gene in regulating the actomyosin cytoskeleton. We tested the hypothesis that TWIST1 regulates *Specc1l* expression during CNCC development and that both factors interact E-Cadherin/β-Catenin adhesion protein complex during CNCC development. *Specc1l* gene has 17 exons plus 5’ and 3’ UTR (Fig. 9A). We mapped two putative regulatory elements in intron 1 and 2 based on epigenetic signatures, including DNase I footprint, H3K27Ac, transcription factor signals, and conservation in vertebrate species (Fig. S10). We performed ChIP-PCR to determine whether TWIST1 *in vivo* binds to E1 and E2 putative enhancer elements. The results showed enrichment of TWIST1 protein at the putative *Specc1l* element E1 for both embryonic time points E9.5 and E10.5. The amount of cross-linked protein to E2 putative enhancer element was increased at E10.5 as indicated by the PCR band intensity but not at E9.5 or E11.5 (Fig. 9B). *Twist1* CKO in CNCCs led to a significant reduction in the mRNA expression of *Specc1l* in tissues extracted from the hindbrain and first pharyngeal arch tissues compared to WT littermate embryos (Fig. 9C). We matched the phenotype in *Twist1* CKO and phospho-incompetent embryos with *Specc1l* loss of function mutant mouse embryos to compare the phenotypic overlap in craniofacial tissues and CNCCs. *Twist1^cko/−^* and phospho-incompetent mutant embryos showed brain hemorrhage, subepidermal blebbing, and severe edema along the neural tube (Fig. 9D, E). Similarly, the *Specc1l^cGT/DC510^* compound mutants of a genetrap (cGT) and a C-terminal 510 amino acid truncation (DC510) alleles (described in Hall et al., 2020) show brain hemorrhage, subepidermal blebbing, and severe edema along the neural tube compared to WT littermate embryos (Fig. 9F). Loss of *Twist1* in CNCCs led to disruption in EMT process and the persistence of cell-cell adhesion in migratory mesenchymal cells (Fig. 9H), and the *in vivo* accumulation of adherens junction proteins β-CATENIN and E-CAD in partially detached cells (Fig. 9J, L) compared to WT mesenchymal cells (Fig. 9G, I, K). Similarly, loss of *Specc1l* function caused the accumulation of membrane-associated β-CATENIN in migratory CNCCs (Fig. 9N-N”) compared to WT CNCCs (Fig. 9M-M”). We tested whether SPECC1L protein interacts with adherens junction proteins using U2OS bone osteosarcoma cell lysate due to high expression in this cell line. SPECC1L and E-CAD were detected in the input of total protein lysate. Furthermore, E-CAD protein was pulled down with SPECC1L in a co-IP assay in a lysate immunoprecipitated by anti-SPECC1L antibodies, and no band was detected in the beads-only negative control (Fig. 9O).

**Fig. 9.**
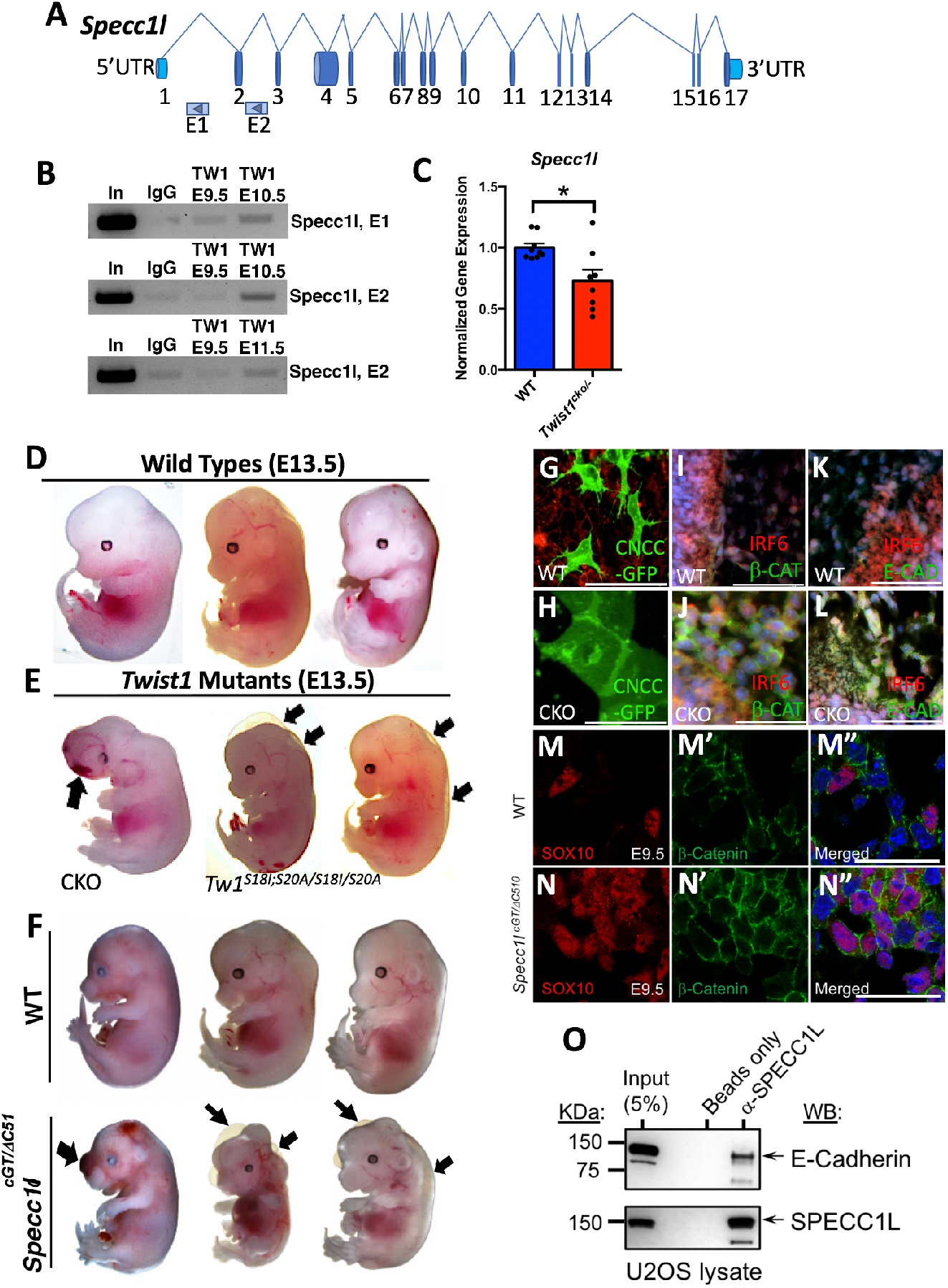
TWIST1 regulates *Specc1l* expression by binding to its putative regulatory elements. (A) Illustration of *Specc1l* gene. (B) By immunoprecipitation, we showed that TWIST1 binds to two *Specc1l* enhancer elements within intron 1 (E1) and 2 (E2) in sorted CNCCs. (C) Measurement of *Specc1l* expression in hindbrain and first pharyngeal arch in WT and *Twist1^cko/−^* littermate embryos. *Specc1l* is significantly decreased in *Twist1^cko/−^* embryos. (D) WT embryos show normal craniofacial development at E13.5, (E) *Twist1^cko/−^* and *Twist1^S18I;20A/S18I;20A^* mutant embryos display craniofacial hemorrhage, subepidermal blebbing, and edema along the neural tube. (F) Images of WT embryos at E12.5 in comparison to *Specc1l^cGT/DC510^* mutant embryo. The *Specc1l^cGT/DC510^* mutant embryos mutant show frontonasal hemorrhage, subepidermal blebbing and edema along the neural tube. (G-H) Confocal images show individual migratory WT CNCCs and a cluster of epithelial-like cells in *Twist1^cko/−^* neural tube explant. (I, J) IF images show no expression of β-CATENIN in mesenchymal cells of WT and a pronounced expression in partially detached mesenchymal cells of *Twist1^cko/−^*. (K, L) No expression of E-CAD in mesenchymal cells of WT was detected and a robust signal in partially detached mesenchymal was observed in cells of *Twist1^cko/−^*. (M-M”) IF images show SOX10 expression and dispersed membrane-associated expression of β-CATENIN in migratory CNCCs of WT embryo. (N-N”) Images show SOX10 expression and pronounced membrane-associated expression of β-CATENIN in migratory CNCCs of *Specc1l^cGT/DC510^* mutant embryo. (O) Co-IP blot in U2OS cells shows the E-CAD is pulled down with SPECC1L protein as shown in the western blot after immunoprecipitation with SPECC1L antibodies. Scale bars represent 50 μm.

### Common variants near *TWIST1* impact human facial shape

The different mouse models of *Twist1* loss of expression and function indicate the crucial role of this gene in regulating the development of orofacial tissues, particularly CNCC-derived structures. To determine the relevance of these animal model findings to humans, we revisited a recently published GWAS meta-analysis of 8246 individuals of recent European ancestry, which identified 203 genome-wide significant signals associated with normal-range facial shape as previously described in White et al., 2021. Moving from minor (A) to major (T) allele of the leading SNP, the major shape changes include a less protrusive and wider nose, a shorter and less protrusive upper lip, and a more prominent chin and forehead. These changes are visible in the morphs and heatmap (Fig. 10A). A signal at 7p21.1 near *TWIST1* was significantly associated facial shape and the lead SNP (rs212672) is located approximately 355 kb downstream of *TWIST1* (Fig. 10B). The signal is shown in the LocusZoom plot in Fig. 10B. When the face was partitioned into global-to-local anatomical segments, the strongest association with the lead SNP was observed for the whole face (p = 1.31E^−20^) and for CNCC-derived facial regions involving the nose and upper lip (see rosette diagram in Fig. 10C).

**Fig. 10.**
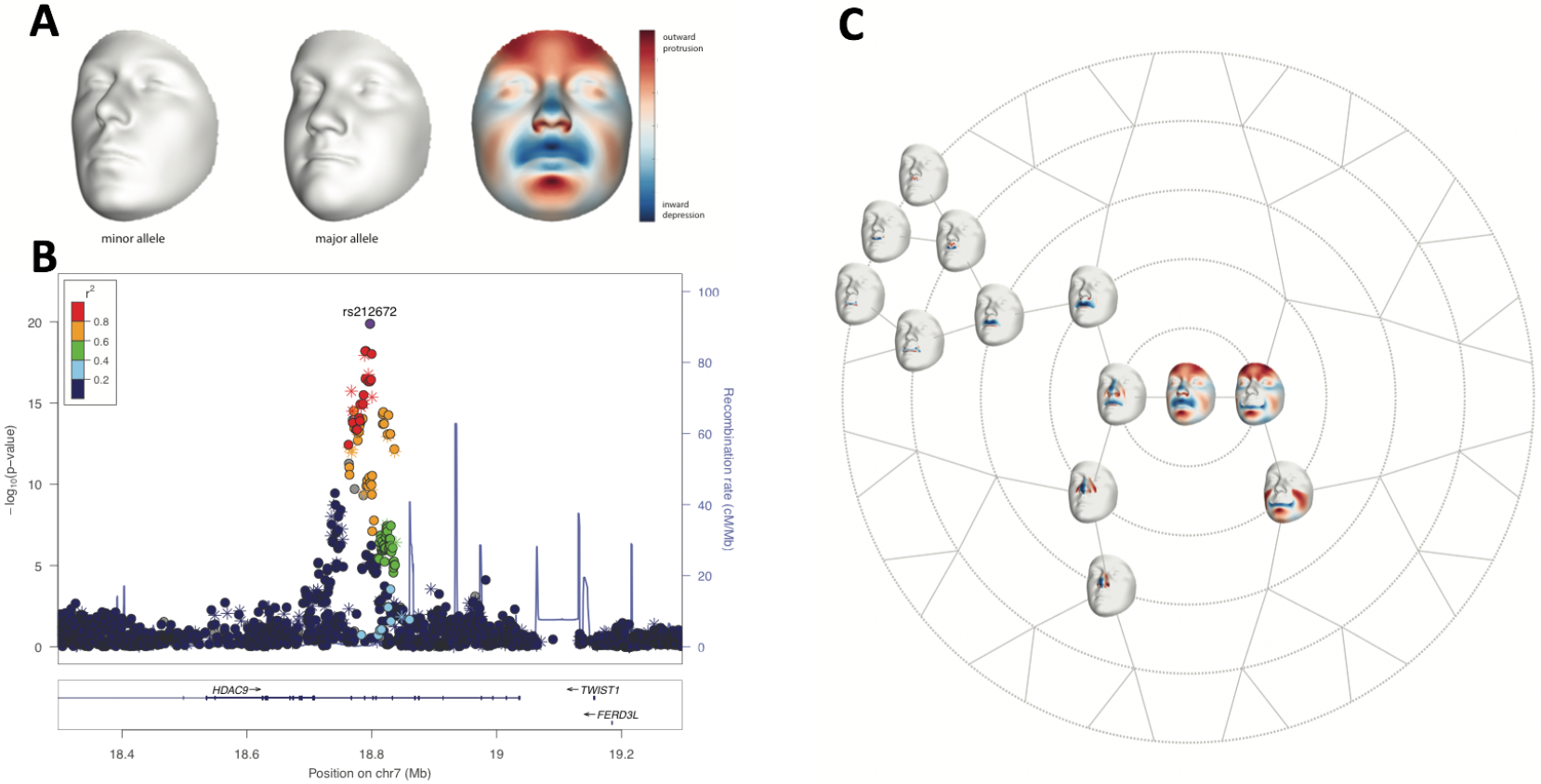
GWAS of 3D human facial surfaces showing a strong signal near *TWIST1* locus. (A) The 3D facial surfaces above the LocusZoom plot show the phenotypic effects of the lead SNP on the whole face, exaggerated in the direction of the minor (A) and major (T) allele SNP variant. The heatmap shows the same effect using the normal displacement in each point comprising the facial surface (∼8000 points) going from the minor to the major allele SNP variant, with blue representing inward depression and red representing outward protrusion. (B) The points in the diagram are color coded based on linkage disequilibrium (*r*^2^) in Europeans with lead SNP rs212672. The asterisks represent genotyped SNPs, the circles represent imputed SNPs. (C) The branching rosette diagram to the right of the LocusZoom plot shows the effect of the lead SNP (as a heatmap) on various facial segments, following unsupervised hierarchical clustering of the full facial surface into increasingly smaller subparts. The nodes in this diagram represented by 3D facial surfaces are the facial segments where the lead SNP reached the nominal genome-wide p-value threshold (p = 5×10^−8^). The heatmaps on each 3D surface show the anatomical boundaries of the segments and can be interpreted as described above. The smallest p-value was observed on the whole face. The other facial segments impacted by the lead SNP involved the nose and upper lip.

## DISCUSSION

Uncovering the function of *Twist1* and *Irf6* in cell fate regulation is crucial to identify associated genes and their regulatory pathways that can control tissue development and differentiation. These associated genes and their upstream regulators might explain the missing heritability in familial craniofacial birth defects. The current study sought to elucidate the spatiotemporal expression and function of *Twist1* and *Irf6* in neural tube and CNCC development. Our work tested the hypothesis that *Twist1* is involved in neural tube development and EMT of CNCCs, while *Irf6* helps define neural tube dorsal edges and structural integrity. Our rigorous IF staining results showed that TWIST1 protein is expressed in an overlapping pattern with tight and adherens junction proteins at the apical side of neural plate and the dorsal edges of neural folds. Furthermore, the *in situ* hybridization data showed that *Twist1* mRNA is weakly expressed in the dorsal edges of the neural folds at E8.5 and E9.5. Although a recent study indicated that few cells of the neural tube express *Twist1* mRNA detected by single cell RNA-seq technology (Soldatov et al., 2019), *Twist1* expression in the neural folds has never been described thus far. Moreover, the scRNA-seq data relies on computational analysis to determine the spatial expression and the relative anatomical distribution of the cell-type population (Soldatov et al., 2019).

Interestingly, *Twist1* expression was detected in round-shape cells residing in the neuroectoderm of the forebrain during the emerging of CNCCs at E8.5 (Fuechtbauer, 1995). To determine the function of TWIST1 apical expression in neural folds, we tested whether TWIST1 interacts with adherens junction proteins. Our Co-IP data showed that TWIST1 interacts with α/β/γ-CATENINS during neural tube closure, and *Twist1* CKO by *Wnt1-Cre* causes disruption of β-CATENIN apical expression which becomes prominently cytosolic. Remarkably, TWIST1 and β-CATENIN have previously been shown to form a complex together with TCF4 in the nucleus (Chang et al., 2015). A recent study demonstrated that conditional knockout of *β-Catenin* by *Wnt1-Cre* causes neural tube closure defects and mutant embryos showed focal loss of N-CADHERIN at the apical side of neural folds (Pieters et al., 2020). This finding is critical because it supports the importance of the apical expression of β-CATENINS and its potential interaction with TWIST1 during neural fold closure.

Previous studies showed that *Twist1* null mice have neural tube defects and disruption in the matrix of the adjacent cephalic mesenchymal cells (Bildsoe et al., 2009; Bildsoe et al., 2013; Chen and Behringer, 1995; Firulli and Conway, 2008). *Twist1* null mice exhibited cranial neural tube closure defects, degeneration of the apical neuroepithelial cells of the neural tube, and increased cell death in migratory CNCCs (Bildsoe et al., 2009; Chen and Behringer, 1995; Ota et al., 2004; Soo et al., 2002). While the activity from the adjacent cephalic mesenchyme contributes to neural fold elevation, it does not explain why the neural folds in *Twist1* null embryos do elevate but fail to bend and close completely. Similarly, in *Twist1* CKO in CNCCs, the neural folds bend as if to join but stop prematurely, preventing the proper closure of the neural tube. Our data also showed that loss of *Twist1* in neuroectodermal cells by the *Wnt1-Cre* or *Wnt1-Cre2* lines cause the formation of ectopic lateral bending points and expansion of the lateral neural folds. These findings may implicate that loss of *Twist1* in the neural folds alters the expression of adherens junction proteins and other epithelial marker genes leading to abnormalities in neural fold bending and closure.

Although the phenotypes caused by a lack of *Twist1* in CNCCs and mesoderm can be explained by an indirect role of *Twist1* to regulate neural tube formation, *Twist1* expression in neural folds can provide a complementary mechanism for explaining the neural tube closure defects in *Twist1* null, and neural tube multiple bending points and expansion in *Twist1* CKO by *Wnt1-Cre*. Although a complete loss of *Twist1* did not disrupt the anterior-posterior patterning of the neural tube at E9.5, there was expansion in a few components of Shh and Fgf pathways in the prospective fore and midbrain (Soo et al., 2002). A direct role in the expansion of these components was not previously stated because *Twist1* expression in neural folds was not described by the *in situ* hybridization assay at E8.5 (Soo et al., 2002). Notably, *Twist1* CKO in mesodermal cells disrupted neural fold elevation and neural tube closure (Bildsoe et al., 2013), and the rescue experiment with WT mesenchymal cells in *Twist1* null chimera embryos indicated that cephalic mesenchymal cells are involved in neural fold elevation (Chen and Behringer, 1995). The previous findings indicated that *Twist1* activity in cranial mesoderm and migratory mesenchymal cells is required for neural fold elevation by regulating the cell-matrix interactions (Bildsoe et al., 2016). This study reports that TWIST1 expression in the neural folds might play a direct role in neural fold bending and closure by interacting with α/β/γ-CATENINS. For the function of IRF6 in neural tube, we have shown that overexpression of *Irf6* in the basal layer of non-neural ectoderm suppresses *Ap2α* in the ectoderm and causes neural tube abnormalities and exencephaly (Kousa et al., 2019). Our current findings show that AP2α expression was expanded into the neural tube in *Irf6* null embryos which could explain the disruption of the integrity of neural tube edges. Also, the glial progenitor cells within the neural tube were disorganized in *Irf6* null embryos with more noticeable gaps between the progenitor cells.

The *Wnt1* ligand is a key regulator in the identification of neural plate border domains and early induction of CNCCs (Parr et al., 1993). The *Wnt1-Cre* transgene drives the expression of *Cre* throughout the neural plate at E8.5 and at later stages during neural tube development (Chang et al., 2015). Therefore, *Wnt1-Cre* was ideal to delete *Twist1* at early stages of its expression if it is present during early neural fold formation. Previous studies showed that *Twist1* CKO in neuroectodermal cells by *Wnt1-Cre* did not disrupt the migration of CNCCs towards the frontonasal and pharyngeal arches. However, the number of migratory *Twist1*-deficient CNCCs was reduced. Based on cell proliferation and apoptotic assays, *Twist1* was considered to act as a survival factor in migratory CNCCs and as a crucial transcription factor for osteoblast lineage-specification (Chen et al., 2007; Chen et al., 2017; Zhang et al., 2012). Apoptosis in *Twist1*-deficient CNCCs might be an end-stage behind the loss of craniofacial bone and cartilage. Yet, no report has indicated whether *Twist1* is involved in the events before CNCC migration and proliferation, or if the migratory *Twist1*-deficient CNCCs are normal mesenchymal cells or are deficient in transition to become multipotent mesenchymal cells. The rationale behind testing these hypotheses is the neural tube phenotype and the loss of craniofacial bone despite the migration of a sizeable amount of CNCCs toward the pharyngeal arches. Furthermore, many cancer studies demonstrated that TWIST1 is a master regulator of EMT (Chen et al., 2017; Zhang et al., 2021), while its function in the EMT process during development is still not well-understood. This study’s data using neural tube explants of *Twist1^cko/−^; Wnt1-Cre*;*R26^dTeG^* mice showed that *Twist1* regulates EMT in CNCCs and a loss of *Twist1* leads to disruption of cell fate transition and cell delamination. Our findings also showed that TWIST1 suppresses the expression of *Irf6* and other epithelial factors during delamination of CNCCs. *Irf6* is a terminal differentiation factor of proliferative epithelial cells in the ectoderm and its expression prevents EMT (Fakhouri et al., 2017; Ingraham et al., 2006; Richardson et al., 2006; Thompson et al., 2019). Notably, the overexpression of *Twist1* in the trunk neural tube converted the pre-EMT neuroectodermal cells into cells that resemble CNCCs (Soldatov et al., 2019). These findings suggest that TWIST1 is involved in cell fate determination of pre-EMT neuroectodermal cells.

Previous findings, including the data from this study, demonstrated that *Twist1^cko/−^* mice have severe craniofacial deformities, including exencephaly, severe mandibular hypoplasia, and a lack of frontonasal and maxillary bone (Bildsoe et al., 2009; Firulli et al., 2007; Vincentz et al., 2008). In humans, haploinsufficiency of *TWIST1* leads to the syndromic form of craniosynostosis and cleft palate (Kunz and Fritz, 1999; Seto et al., 2007; Takenouchi et al., 2018). It has been shown that *Twist1*-deficient mesenchymal cells do not have mesenchymal characteristics as observed in mouse chimeras created from wild type embryos injected with M-twist deficient embryonic stem cells (Chen and Behringer, 1995), and that *Twist1* is essential for CNCC survival to give rise to CNCC-derived craniofacial formation as shown by an increased level of mesenchymal apoptosis in *Twist1^cko/−^* (Bildsoe et al., 2013; Chen et al., 2007; Zhang et al., 2012). Although a cause for the cell death observed in *Twist1* CKO in CNCCs was not determined, our current findings suggest that the *Twist1*-deficient CNCCs did not complete their EMT transition and maintained the expression of differentiation factors. Therefore, they possibly did not become potent cells that can respond to extrinsic stimuli. Recent studies on the EMT process showed that a complete conversion from an epithelial to a mesenchymal state is not necessary for cancer cells to migrate, and multiple metastable states during cell transition were also recently described in cancer studies (Cousin, 2017; Nieto et al., 2016; Thiery et al., 2009). However, we have not observed a collective cell migration or epithelial-like cells in WT neural tube explants or even *in vivo* histological staining.

Determining the *in vivo* role of TWIST1 post-translational modification is vital in delineating its role in neural tube and CNCCs. TWIST1 is a phospho-protein and multiple phospho-residues are important for its stability, nuclear translocation, and transcriptional regulation of target genes as per previous cancer studies (Bourguignon et al., 2010; Zhao et al., 2017; Xu et al., 2017). *Twist1* phosphorylation increases tumor cell motility in squamous cell carcinoma of the head and neck (Alexander et al., 2006; Su et al., 2011). Overexpression of c-Src kinase also increased TWIST1 phosphorylation leading to nuclear localization in breast tumor cells (Bourguignon et al., 2010). In addition, overexpression of *Twist1* phosphomimetic T125D; S127D led to limb abnormalities in transient transgenic mouse embryos (Firulli et al., 2007). Notably, the phospho-regulation of the TQS (T125;Q126;S127) motif in the helix 1 domain of *Twist1* has been shown to modulate the dimerization with other transcription factor partners *in vitro* (Firulli et al., 2005). Our *in vivo* data showed that TWIST1 is mostly cytoplasmic in some apical cells of the neural folds and becomes restricted to the nucleus of detached migratory CNCCs toward the pharyngeal arches. We suggest that post-translational phosphorylation of TWIST1 is crucial for regulating its cellular localization and activity in development. The *Twist1^S18;20A/S18;20A^* and *Twist1^S68A/S68A^* mutants embryos showed several craniofacial abnormalities, including epidermal blebbing, severe edema along the neural tube, meningeal detachment, and neural tube defects. The skeletal staining of these phospho-incompetent mutant embryos showed bone loss and a reduction in skull mineralization of the frontonasal and maxillary bone. Yet, the *in vivo* function of the two phospho-sites need to be delineated, including their importance in protein stability, nuclear translocation, EMT process, cell survival, and cell migration. The data of this study describes a new role of *Twist1* involved in the EMT of CNCC during embryogenesis.

The data generated on the differentially expressed genes in *Twist1^cko/−^* embryos demonstrated that many factors involved in intercellular adhesion proteins and cytoskeletal remodeling are significantly altered in the hindbrain and first pharyngeal arch. Previous studies showed that mutations in adhesion protein and cytoskeletal regulator genes could cause neural tube closure defects and improper delamination of CNCCs, leading to craniofacial disorders (Wilson et al., 2016). In addition, EMT induced by TWIST1 promotes the formation of tubulin-based microtentacles that penetrate the junctions between endothelial layers (Whipple et al., 2010). One of the genes contributing to this delamination defect in mice is *Specc1l* (Wilson et al., 2016). Both *Twist1* and *Specc1l* are involved in NC cell development and impact craniofacial structures in mice and humans. Mutations in TWIST1 and SPECC1L lead to craniosynostosis and orofacial clefting in humans (Bai et al., 2021; Bhoj et al., 2019; Hall et al., 2020). *Specc1l* is a gene that encodes a novel coiled-coil domain containing protein, which stabilizes microtubules and actin of the cytoskeleton (Bhoj et al., 2019; Gfrerer et al., 2014). SPECC1L colocalizes with both actin and tubulin (Saadi et al., 2011; Wilson et al., 2016). Thus, without sufficient SPECC1L, actin-cytoskeleton reorganization and cell adhesion are significantly affected (Saadi et al., 2011). The overlap in cellular function between *Twist1* and *Specc1l* and the remarkable phenotypic overlap in *Twist1* phospho-incompetent and *Specc1l* mutant mouse embryos, including the craniofacial abnormalities due to possibly the accumulation of cell adhesion factors in migratory mutant CNCCs, suggesting that both genes are involved in a similar regulatory pathway to control cytoskeleton reorganization during cell delamination and migration. This study shows that TWIST1 directly binds to putative regulatory elements in the intronic regions of *Specc1l* gene and loss of *Twist1* led to a significant reduction of *Specc1l* in hindbrain and first pharyngeal arch tissues.

In conclusion, our findings emphasize the conservation of TWIST1 function in regulating the EMT process in CNCCs by controlling the expression of epithelial genes, including *Irf6*, intercellular adhesion proteins, and cytoskeletal regulators. TWIST1 interacts with CATENINS that are apically expressed in the dorsal side of neural tube, and also regulates *Specc1l* expression by directly binding to its putative intronic regulatory elements. Furthermore, *Specc1l* and *Twist1* play a similar role in the delamination of cranial neural crest cells by remodeling the cell-cell adhesion and cytoskeletal reorganization. Thus, previous reports and the current study both provide a plausible explanation for the overlapping phenotypes in mice and humans due to loss of TWIST1 function and/or expression by identifying direct target genes involved in cell adhesion and cytoskeletal regulation. Moreover, as our results highlight, reanalysis of the GWAS data in humans also shows that variants near *TWIST1* impact normal-range facial shape, including CNCC-derived structures of the midface (White et al., 2021). Interestingly, the most significant SNP near *TWIST1* overlaps with epigenetic enhancer marks in which the elements with the epigenetic marks were tested for enhancer activities to recapitulate TWIST1 endogenous expression (Hirsch et al., 2018). Understanding the transcriptional regulation of *TWIST1* and its function in controlling the associated-regulatory pathway involved in cell fate determination in neural tube and CNCCs can lead to the identification of the missing heritability in families that are affected and at high risk for craniofacial birth defects due to damaging mutations in IRF*6*, *TWIST1,* or their regulatory elements and target genes.

## MATERIALS AND METHODS

### Mouse strains

All the mice used in this study were generated using the C57BL/6J genetic background. *Twist1* heterozygous mice were generated using *EIIA-Cre* mouse line using the B6;129S7-*Twist1^fl/fl^* (obtained from MMRRC supported by NIH). The mouse lines 129S4.Cg-*E2f1^Tg(Wnt1-cre2)Sor^*/J (catalog #: 022137), BL6.129(Cg)-*Gt(ROSA)26Sor^tm4(ACTB-tdTomato,-EGFP)Luo^*/J (catalog#: 007676), and BL6.129S4-*Gt(ROSA)26Sor^tm1Sor^*/J (catalog #: 003474) were obtained from the Jackson Laboratory. The two different cell tracing strategies R26 *^tm4(ACTB-tdTomato,-EGFP)^* and R26 *^tm1.lacZ^* were used to track *in vivo* CNCC formation and migration. *Twist1^S18;20A/+^*, *Twist1^S42A/+^*, *Twist1^S68A/+^*, and *Twist1^T121;S123A/+^* phospho-incompetent founders were generated using CRISPR/Cas9 methodology at the Baylor College of Medicine and the founders were backcrossed to wild type C57BL/J mice for 6 generations. Genomic DNA from the founders was sequenced to confirm the substitution of S18/20 and S68. *Twist1^S18I;20A/+^* and *Twist1^S68A/+^* heterozygous mice were crossed to obtain homozygous *Twist1^S18I;20A/S18I;20A^* and homozygous *Twist1^S68A/S68A^* embryos. These embryos were generated to determine the effects on the CNCC-derived craniofacial bone and cartilage and the earliest time-point of detecting a craniofacial pathology. The animal work was approved by the CLAMC committee at UTHealth Houston under the Approved Animal protocol # AWC-19-0045.

### Mouse handling, embryo extraction, and genotyping

Embryos were extracted based on the presence of a copulation plug and the number of days after the last known delivery. Pregnant females were euthanized with CO_2_ followed by cervical dislocation. Embryos were genotyped for *Twist1^fl/fl^*, *Twist1^+/−^*, *Twist1^−/−^*, *Twist1^fl/−^*, *Wnt1-Cre1, Wnt1-Cre2*, *Irf6^−/+^, Twist1^S18I;20A/S18I;20A^*, and *Twist1^S68A/S68A^* alleles via DNA extraction, allele-specific PCR, and gel electrophoresis.

### Histological and immunofluorescent staining

Embryos of WT, *Twist1^−/−^*, *Twist1^cko/−^*, and phospho-incompetent lines were collected for histological and IF staining. Immunofluorescent staining was performed as previously described (Fakhouri et al., 2012). Briefly, mouse tissues were deparaffinized and rehydrated in a series of ethanol dilutions. The slides were boiled for 10 minutes in 10mM sodium citrate buffer for antigen retrieval. Sections were blocked with anti-mouse IgG Fab fragment for 30 minutes, and then with 10% (v/v) normal goat serum and 1% BSA (v/v) in PBS for 1hour, then incubated overnight at 4**°**C with the following primary antibodies: AKT1/2/3 (1-150, sc-81434,), β-CATENIN (1-200, sc-7963), CLAUDIN1 (1-200, sc-166338), N-CADHERIN (1-200, sc-59987,), OCCLUDIN (1-150, sc-133256), and VIMENTIN (sc-6260), E-CADHERIN (1-150, BD-610182), mouse anti-TWIST1 (1-200, ab50887), SOX9 (1-200, ab185966), mouse anti-ZO-1 (sc-33725), mouse anti-TFAP2α (PCRP-TFAP2A-2C2, DSHB, Iowa), rabbit anti-CD73 (1-150, ab133582), rabbit anti-CD206 (1-150, ab64693), Phalloidin (1-1000, ab176757), and rabbit anti-IRF6 antibodies (1-500, Fakhouri et al. 2012). The secondary antibodies were goat anti-rabbit (A21429, Molecular Probes, CA) and goat anti-mouse (A11029, Molecular Probes, CA). We marked nuclei with DAPI (D3571, Invitrogen, CA). An X-Cite Series 120Q laser and a CoolSnap HQ2 photometric camera (Andor Neo/Zyla) installed in a fluorescent microscope (Nikon Eclipse Ni) were used to capture images. For the IF staining, sections from five embryos of each genotype, WT and *Twist1^cko/−^*, were used for each tested embryonic time point.

### *Twist1* mRNA probe for *in situ* hybridization

The *Twist1* probe for *in situ* hybridization was designed to recognize a 0.8 kb region in the 3’ UTR region of mouse *Twist1*, starting at 188 bp and ending at 1062 bp after the *Twist1* coding sequence ends. This avoids any conserved regions homologous with other bHLH factors and, encompasses the sequence of the shorter probe described in Wolf et al., 1990. Further details are described in Fig. S2.

### Whole mount embryo β-Gal staining

*R26^Tm1^* transgenic mice were used to perform X-gal staining for cell tracing in whole-mount embryos. Whole-mount embryo staining at E9.5, E11.5, and E15.5 was performed as previously described (Metwalli et al., 2018). Stained wild type and *Twist1^cko/−^* embryos were then visualized and analyzed compared to wild-type embryos using a stereomicroscope and NIS Elements AR software. The X-gal staining was performed on more than three biological replicates of each genotype and tested embryonic stage.

### Skeletal staining

*Twist1* phospho-incompetent embryos at E16.5 were dissolved in 2% KOH until the embryo’s skin dissolves, and then stained with Alcian blue solution overnight for cartilage and Alizarin red for bone as previously described (Thompson et al., 2019). After that, the embryos were submerged in 10% glycerol and 1% KOH to remove excess stain and remaining tissues for 3-5 days.

### Neural tube explant excision and CNCC culture

E8.5-E9.5 embryos from the *Twist1^fl/−^* ;*Irf6^−/+^*;*Wnt1-Cre2* crosses described previously were extracted from their amniotic sacs and maintained in an organ culture medium. Using microdissection instruments, E8.5-9.5 embryos were removed from their amniotic sacs. Transverse cuts were made through the optic vesicle and first pharyngeal arch. After the removal of extraneous tissue, the mid and hindbrain regions were sectioned and cultured on Collagen Type I-coated petri dishes (10µg/cm^2^) in alpha-MEM medium (10% fetal bovine serum, 100 U/mL penicillin G, 100 μg/mL streptomycin, 1% non-essential and essential amino acids) and placed in a 5% CO_2_ incubator at 37°C overnight to allow for appropriate adherence.

### Time-lapse image acquisition and analysis

Time-lapse imaging of explants was performed using the A1R MP Confocal and Multiphoton Microscope (Nikon Instruments). Using overlaid contrast of red (TRITC), green (FITC), and blue (Hoechst) channels, fluorescence color changes from tissue explants were analyzed. Green fluorescence indicated the GFP reporter found only in CNCCs, while red fluorescence indicated all other tissues that express RFP, with Hoechst serving as a nuclei counterstain. Neural tube explants were imaged in an environmentally controlled chamber maintained at 37°C and 5% CO_2_, and the migration of CNCCs was captured using automated acquisition over a 21 hour period. The net migration of cell bodies over a 10-hour period was calculated using IMARIS software (Bitplane AG, Nikon). Parameters included a spot diameter and connected components algorithm used for cell tracking. Cells also were tracked frame-by-frame using the manual-tracking spot feature of IMARIS 9.2.1. About 16 WT and 22 *Twist1^cko/−^* CNCCs were tracked. A two-tailed Mann–Whitney U-test was used with a significance level of ****p < 0.0001.

### Western blot, protein phosphorylation, and RTqPCR

We performed immunoblots to measure the quantitative levels of TWIST1 (ab50887), SNAIL2 (sc-166476), AKT1 (sc-81434), E-CAD (sc-8426), N-CAD (sc-59987), β-CATENIN (sc-7963), and WNT3 (sc-74537). We also measured the mRNA level of these genes in addition to *Pi3k, Ck2, Erk1, Vimentin, Occludin, Msx1, Tfap2α, Hand2, Yap, Wnt1, RhoA, RhoC, Arhgap29, Sox2, Sox10, HoxD10, Jag1, c-Src*, and *mTor*. The levels of phosphorylated and unphosphorylated forms of TWIST1 at S42, S68, and S125 were detected via immunoblot as previous described (Fakhouri et al., 2017). We received aliquots of rabbit polyclonal antibodies for TWIST1 P-S42 and P-S127 from Dr. Brian A. Hemmings at the Friedrich Miescher Institute for Biomedical Research, Basel (Vichalkovski et al., 2010). We purchased the monoclonal TWIST1 phospho-S68 from Abcam (ab187008). Monoclonal antibodies for TWIST1 (ab50887) were used for IF and immunoblot, and Co-IP. We used the same antibodies of E-CAD and β-CATENIN for IF, western and Co-IP. For the digestion with phosphatase enzyme, Protein Phosphatase 2A1 bovine (Millipore-Sigma, P6993) was used to digest 100 μl total protein extracted from O9-1 CNCCs. About 10μl PP2A in 50mM TRIS-HCl buffer containing 2-mercaptoethanol was used and the solution was incubated at 30 ^ο^C for 30 minutes and then was analyzed by western blot using TWIST1 monoclonal antibodies (ab50887).

### Sanger Sequencing

Purified genomic DNA from mouse tail snips and purified PCR products were submitted for sanger sequencing at Genewiz Company.

### Co-IP assay for protein-protein interactions

We extracted total protein from mid and hindbrain tissues dissected from embryos at E9.5. Tissues were ground with a plastic pestle on dry ice and the samples were frozen and thawed twice before centrifugation to remove undissolved materials. Purified total protein was incubated with Protein A/G conjugated to magnetic beads. We used monoclonal antibodies (Abs) for TWIST1 (ab50887), E-CAD (sc-8426), and polyclonal Abs for β-CATENIN that can also detect α/γ-CATENINS (sc-7963).

### ChIP-PCR for TWIST1 binding to *Specc1l* putative regulatory elements

Chromatin immunoprecipitation (ChIP) was performed in tissues dissected from hindbrain and first pharyngeal arch. We used TWIST1 monoclonal antibodies to detect the binding to two putative enhancer elements within intron 1 and 2 of Specc1l gene. We also pulled down fragmented chromatins with IgG antibodies as negative control for non-specific binding. The ChIP-PCR was performed as previously described (Kouse et al., 2019).

## Acknowledgements

We thank April Zhang and Yazan Hasan for their excellent help with the immunofluorescent imaging and histological staining of the mouse embryonic tissues. We are very grateful to Dr. Brian A. Hemmings at the Friedrich Miescher Institute for Biomedical Research, Basel, who sent us sufficient aliquots of rabbit polyclonal antibodies for TWIST1 P-S42 and P-S127. We thank Dr. Brendan Lee and Ms. Racel Cela from Baylor College of Medicine for their tremendous help maintaining the *Twist1* phospho-incompetent mouse founders at the Baylor mouse facility. We further thank the Genetically Engineered Mouse Core at Baylor College of Medicine for generating the Twist1 phospho-incompetent mouse lines.

---

**Supplementary Table 1.**
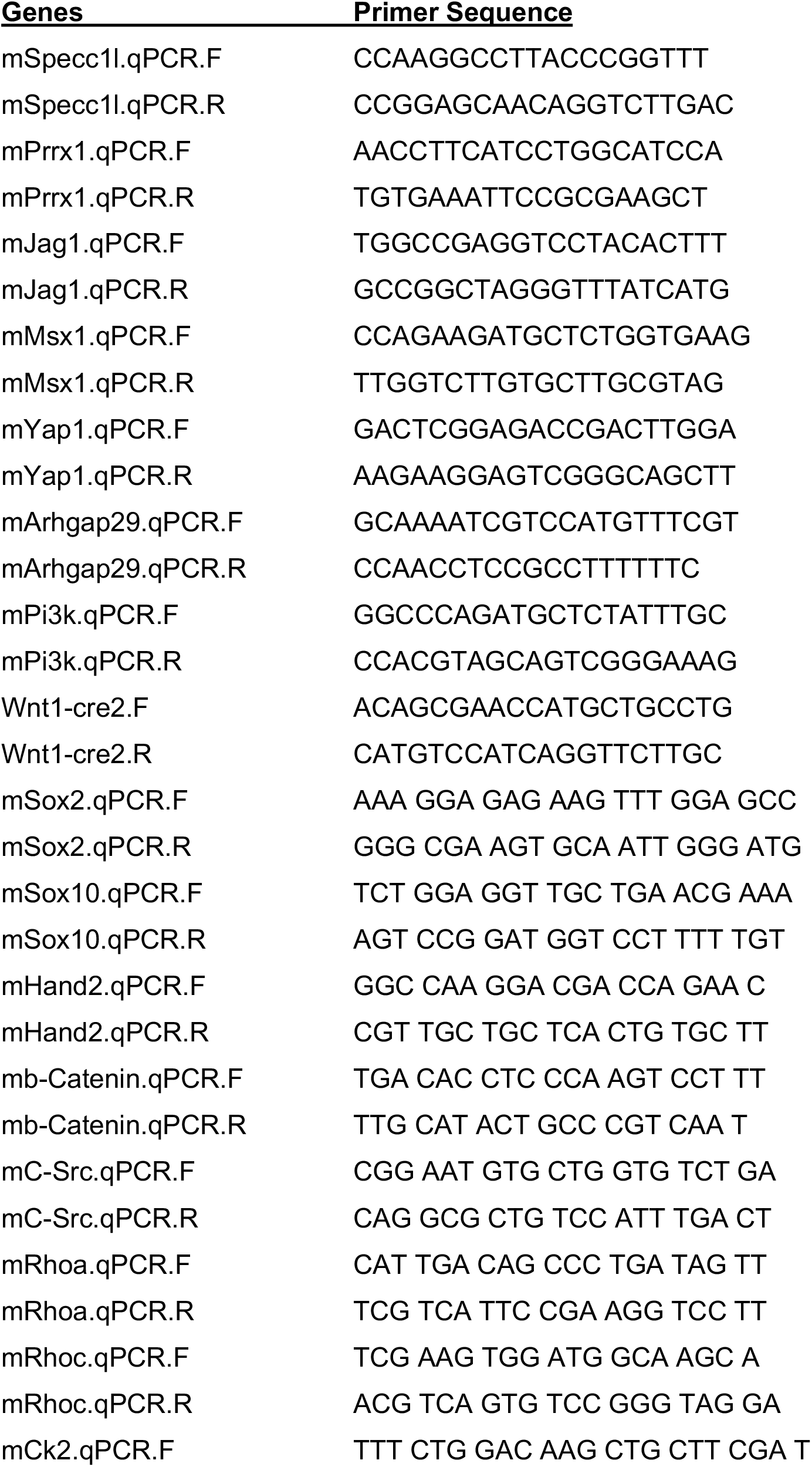
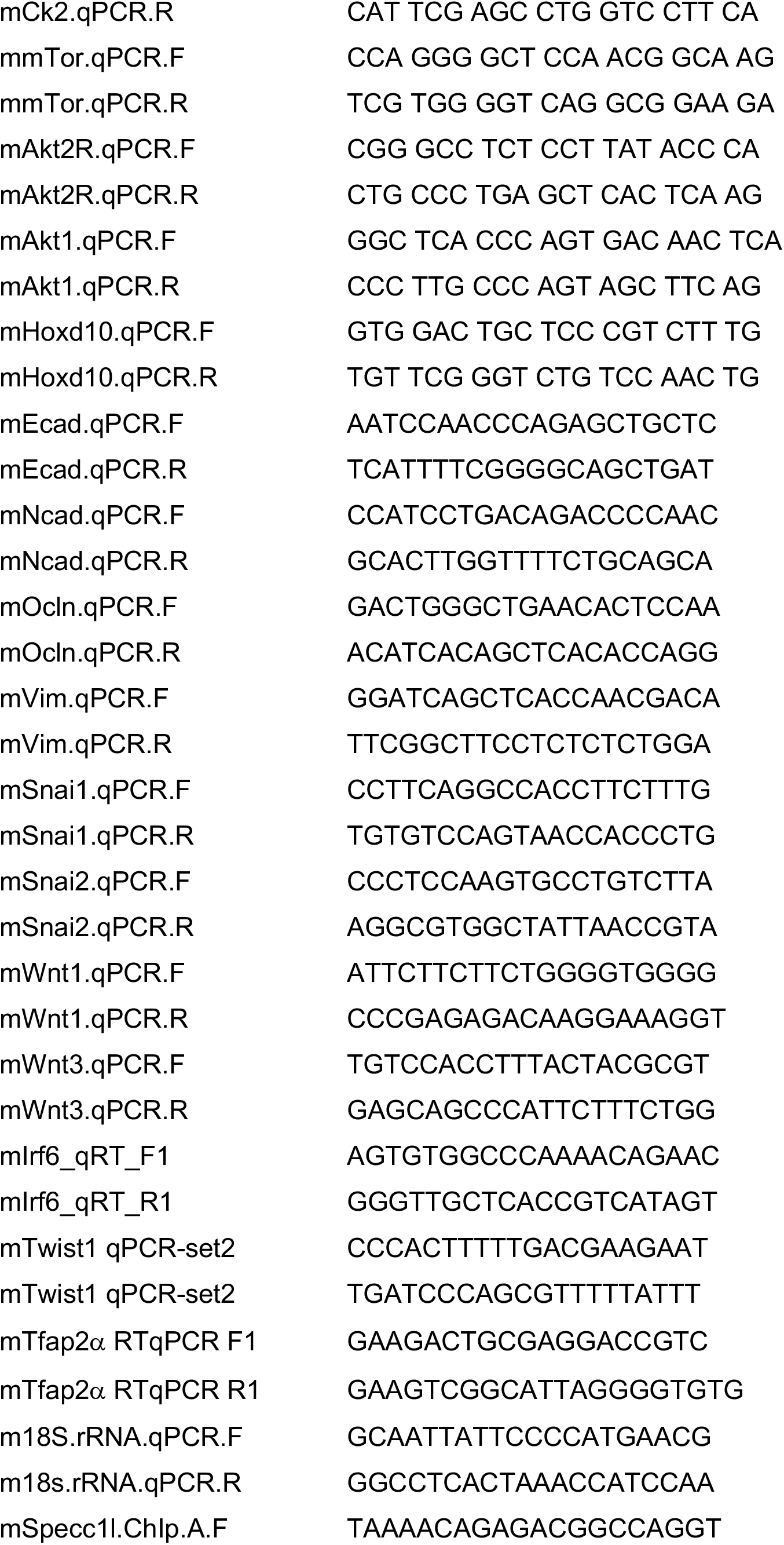
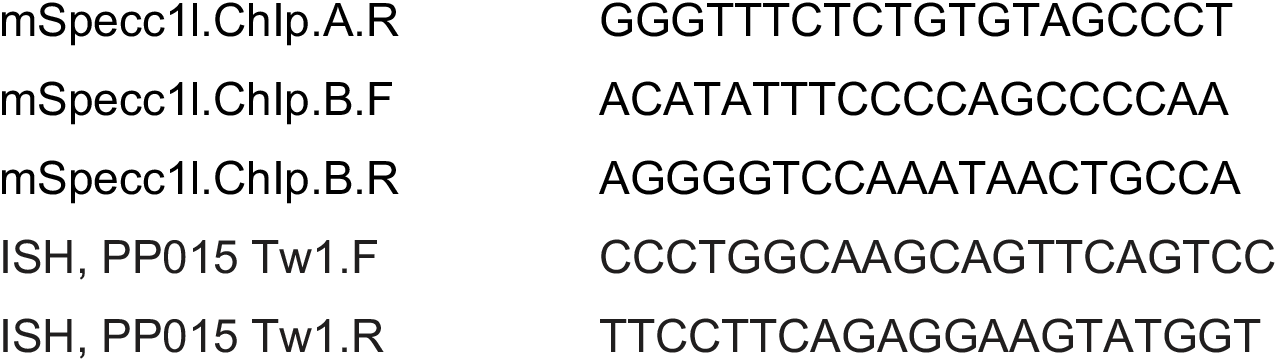
A list of genes and primers was used to quantify mRNA by RTqPCR, chromatin signal by ChIP-PCR, and cellular mRNA by *in situ* hybridization.

**Supplementary Fig. 1.**
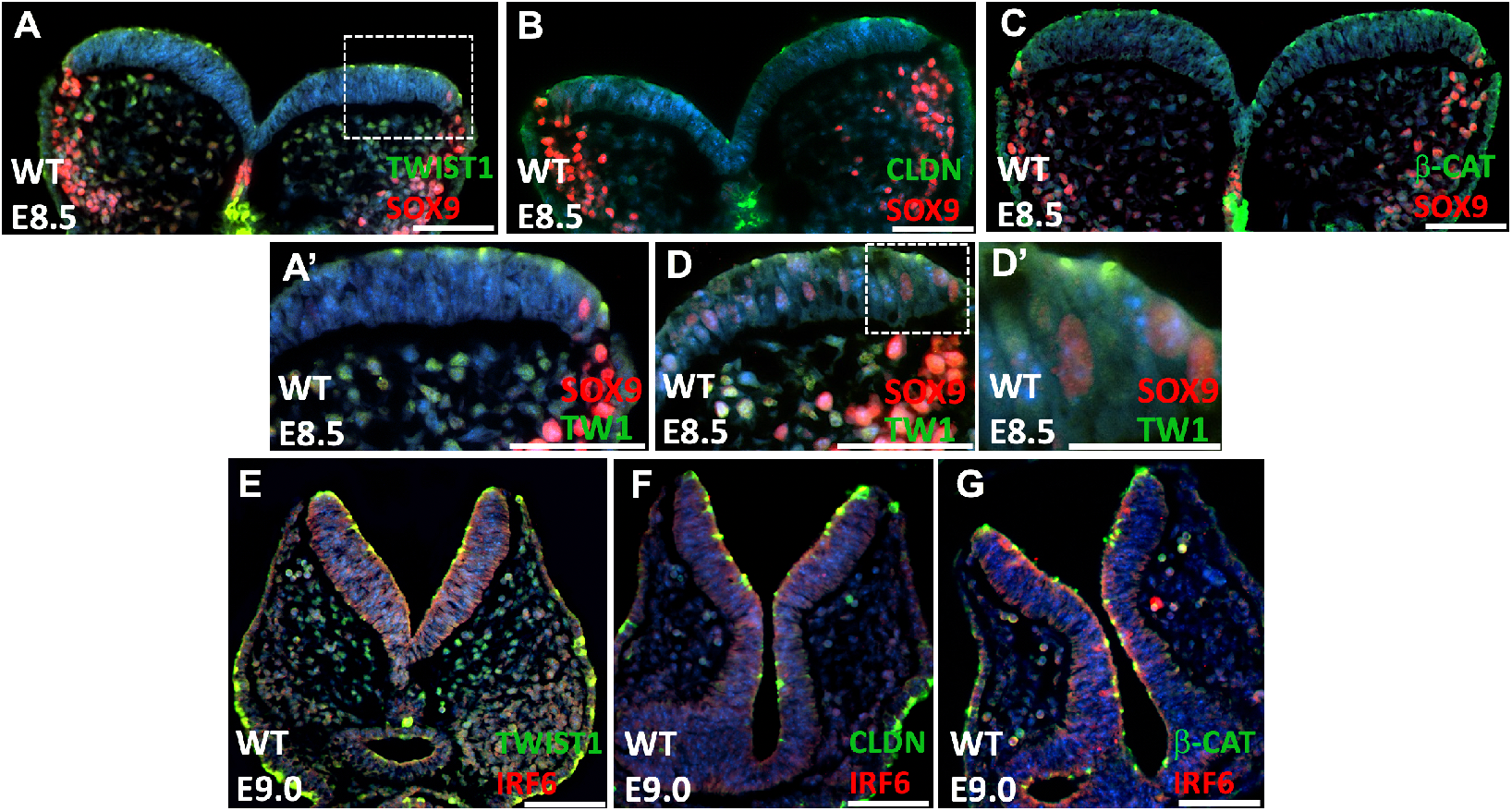
Immunofluorescent staining of TWIST1, SOX9, CLAUDIN, IRF6, and β-CATENIN. (A-C) TWIST1 is apically expressed at the dorsal side of the neural plate, similar to CLAUDIN and β-CATENIN. (D, D’) TWIST1 is also expressed in the cells residing at the neural plate borders and partially overlaps with SOX9 expression at E8.5. (E) TWIST1 is apically expressed in neural folds and at dorsal edges of neural folds. (F, G) Expression of CLAUDIN and β-CATENIN at the apical side of the neural folds at E9.0.

**Supplementary Fig. 2.**
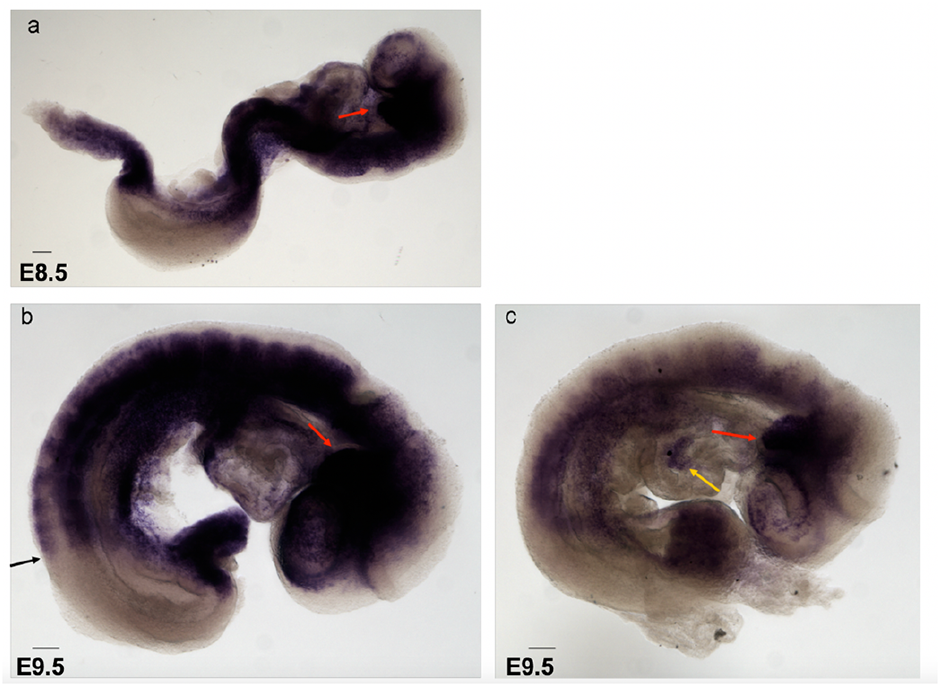
Whole mount *in situ* hybridization for *Twist1* mRNA at E8.5 and E9.5 embryos. The positive staining is depicted in dark purple color in craniofacial, trunk and limb regions. To produce the RNA probe, BAC was extracted from clone bMQ-350m18 (GeneService). PCR primers were designed to amplify a 0.8 kb fragment in the 3’ UTR region of *Twist1* gene. The fragment was amplified, subcloned into TOPO, and then sequenced using the M13 primers. Sequencing revealed that clone WOTO2 contained a correct insert which was used to produce a RNA probe complementary to *Twist1* mRNA using the T7 promoter site for transcription. The linearized and purified DNA plasmid was used as a template for T7 RNA polymerase (Roche, 10881767001) and the probe was labeled with the DIG-UTP for the *in situ* hybridization.

**Supplementary Fig. 3.**
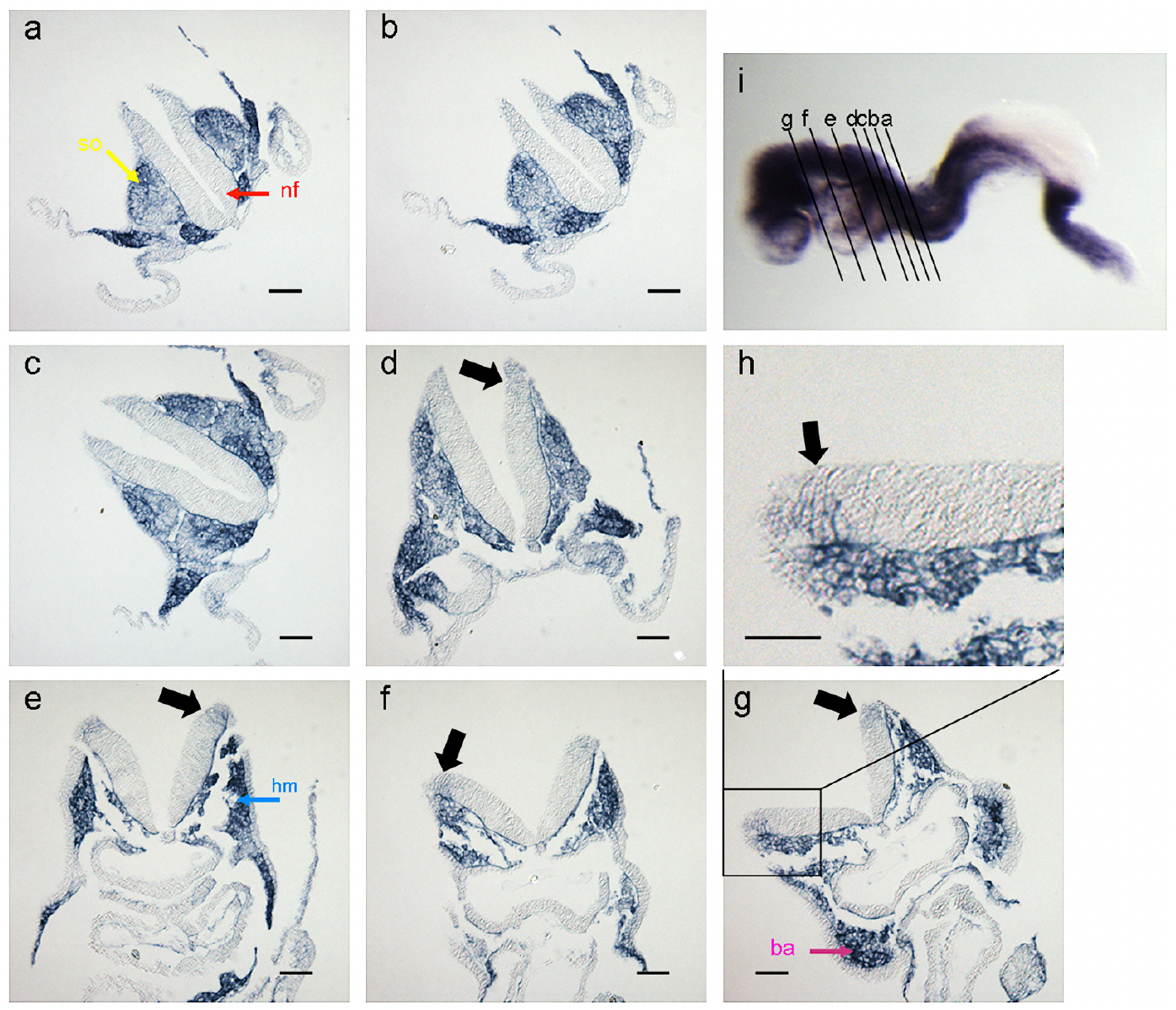
*Twist1* is weakly expressed in the dorsal neural plate at E8.5. Whole mount *in situ* hybridization for *Twist1* of an E8.5 embryo sectioned from posterior to anterior (a to g) as shown in (i). Magnification of the box in (g) focusing on the dorsal neural tube (h). Neural fold (nf, red arrow), somite (so, yellow arrow), head mesenchyme (hm, blue arrow), branchial arch (ba, purple arrow). Scale bars represent 50 μm.

**Supplementary Fig. 4.**
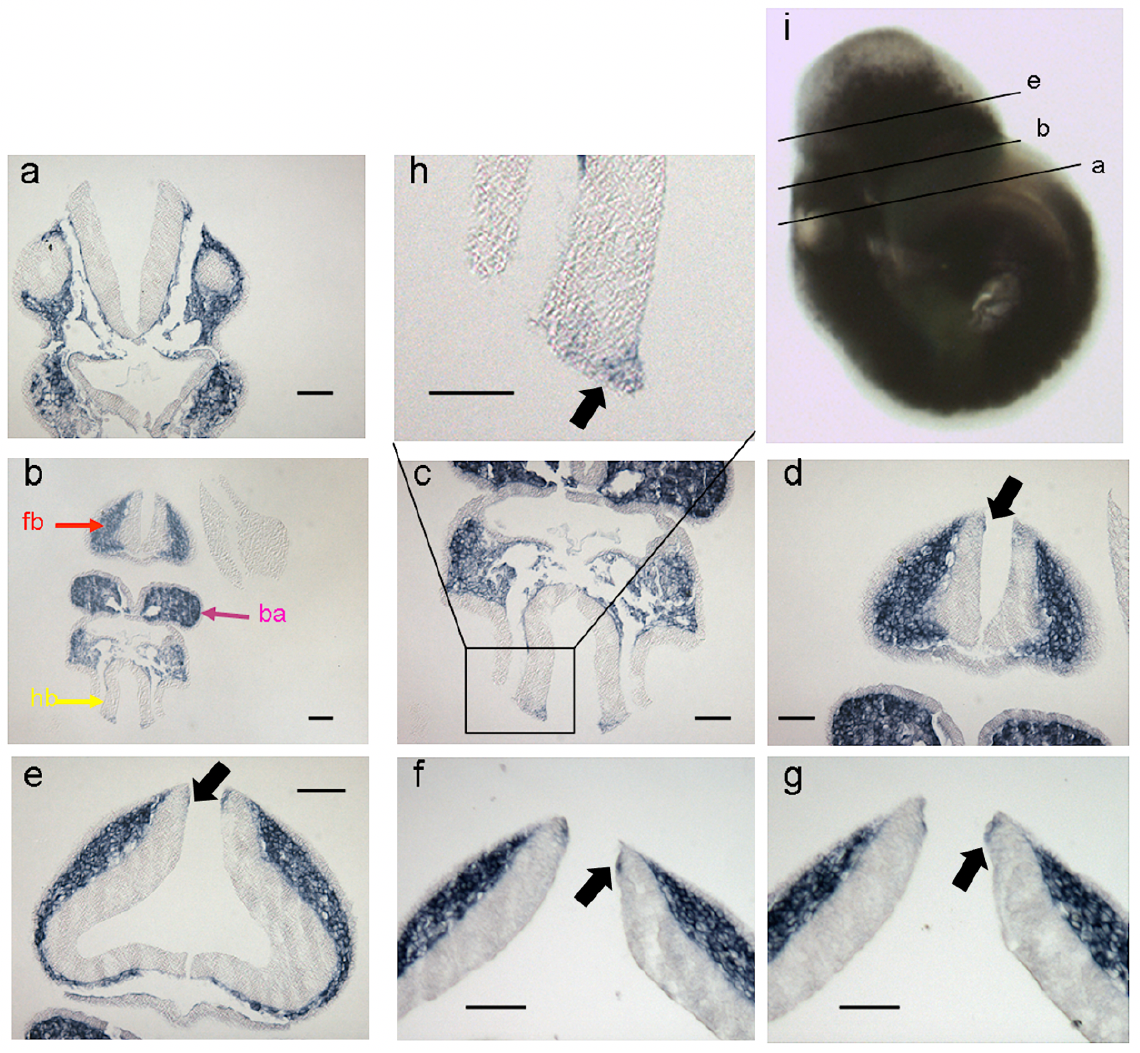
*Twist1* is weakly expressed in the dorsal neural fold at E9.5. Whole mount *in situ* hybridization for *Twist1* of an E9.5 embryo sectioned from posterior to anterior (a to g) as shown in (i). Magnification of the box in (g) focusing on the dorsal neural tube. Forebrain (fb, red arrow), hindbrain (so, yellow arrow), branchial arch (ba, purple arrow). Scale bars represent 50 μm.

**Supplementary Fig. 5.**
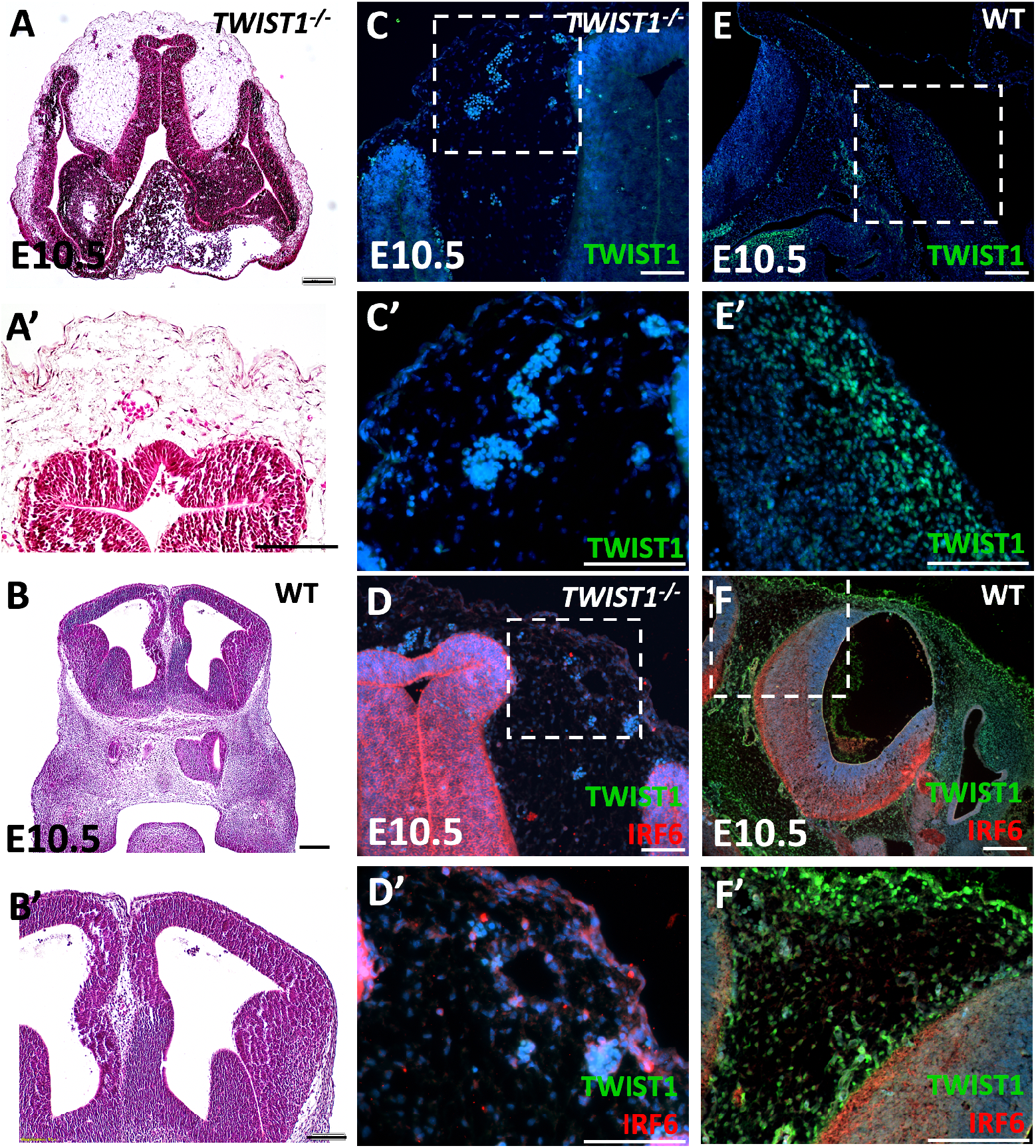
Validation of anti-TWIST1 antibodies in Twist1 null embryonic tissues. (A, A’) Histological staining of coronal section of *Twist1* null embryo showing the morphology of the frontal neural tube and complete loss of nasal and oral tissues at E10.5. (B, B’) Histological staining of the coronal/transverse section of wild-type embryo shows the frontal neural tube morphology. (C, C’) TWIST1 is not detected in *Twist1* null embryo using monoclonal anti-TWIST1 antibody by IF staining. (D, D’) IRF6 is highly expressed in *Twist1* null embryo using dual staining for TWIST1 and IRF6. (E, E’) TWIST1 is detected in migratory mesenchymal cells in comparable tissues of wild-type embryos using the same monoclonal antibody and conditions (F, F’) IRF6 and TWIST1 are highly expressed in wild-type embryos using the same monoclonal antibody dual staining for TWIST1 and IRF6. Scale bars represent 100 μm.

**Supplementary Fig. 6.**
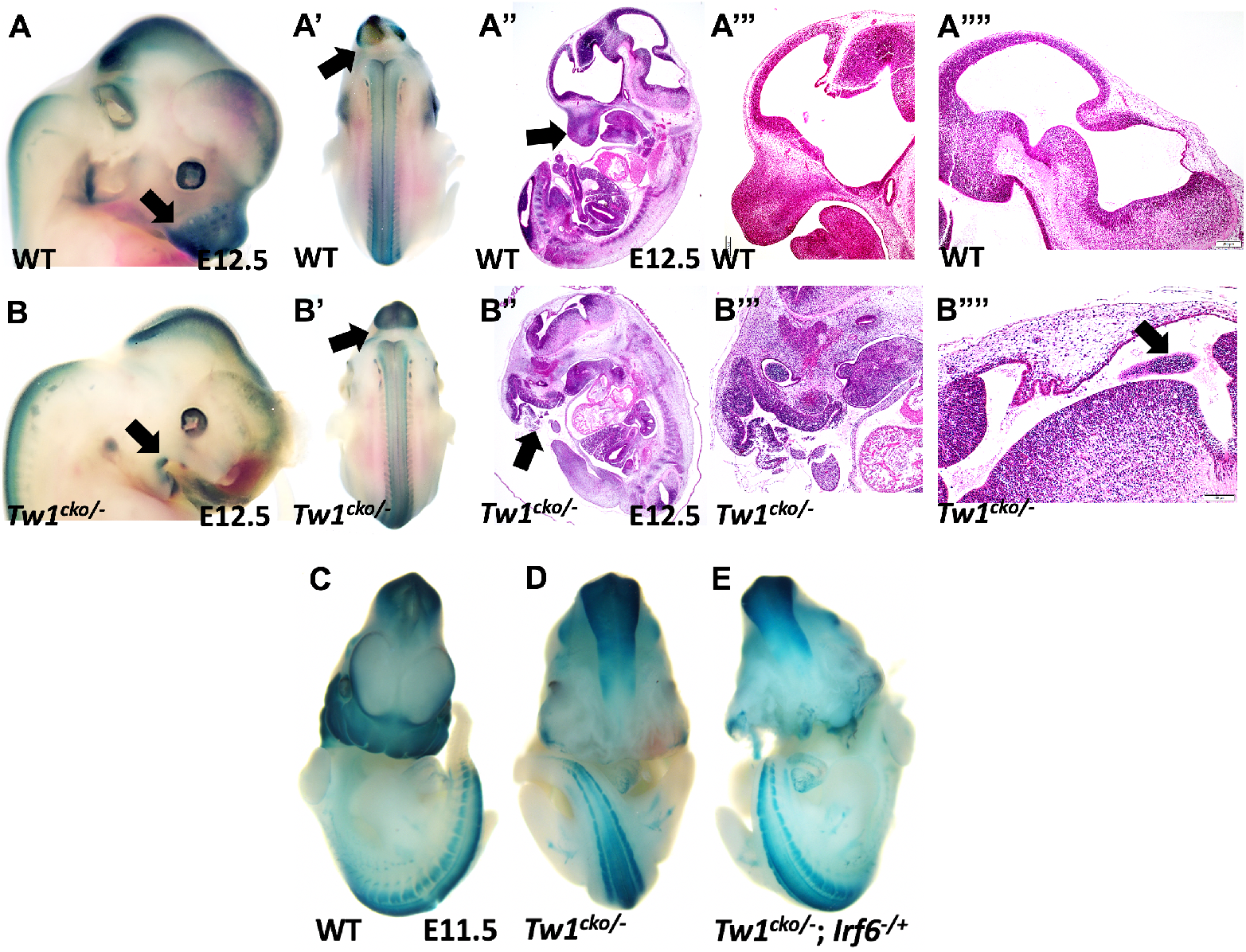
Whole mount X-gal and histological sections were staining in wild type and *Twist1^cko/−^* mouse embryos at E12.5. (A, A’) Whole mount X-gal of WT shows blue staining in frontonasal, jaw processes, midbrain, and trunk neural tube. (A”-A””) Histological staining shows normal brain development in wild type. (B, B’) Whole mount X-gal of WT shows blue staining in frontonasal, jaw processes, midbrain, and trunk neural tube. (B”-B””) and severe brain abnormalities in *Twist1^cko/−^*mouse embryos. (C-E) Whole mount X-gal staining of wild type, *Twist1^cko/−^*, and *Twist1^cko/−^*; *Irf6^−/+^* embryos at E11.5. Compound heterozygous embryo shows severe frontonasal deformities.

**Supplementary Fig. 7.**
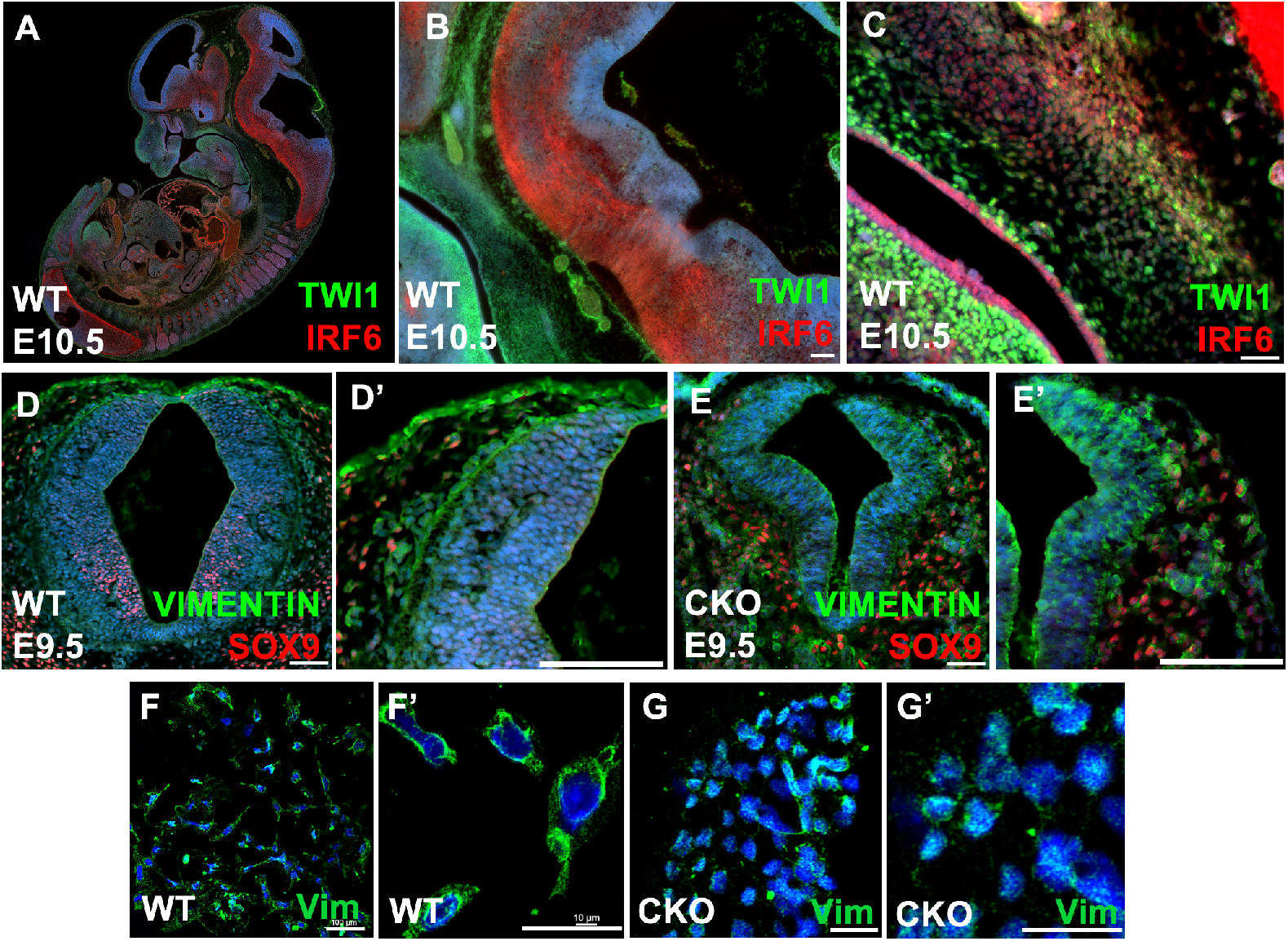
Immunofluorescent staining of IRF6, TWIST1, SOX9, and VIEMENTIN in the neural tube and CNCCs. (A-C) IRF6 is highly expressed in neural tube and oral epithelium, while TWIST1 is highly expressed in migratory mesenchymal cells along the neural tube and in frontonasal and pharyngeal arches. (D-D’) SOX9 is expressed in neuroectodermal progenitors of glial cells in neural tube, while VIEMNTIN is detected in migratory mesenchymal cells along the neural tube. (E, E’) SOX9 was not observed in neural tube and VIEMNTIN was highly expressed in neural tube and partially detached cells along the neural tube. (F-G’) VIEMNTIN is expressed in migratory CNCC emerged from neural tube explants of wild type and *Twist1^cko/−^* embryos.

**Supplementary Fig. 8.**
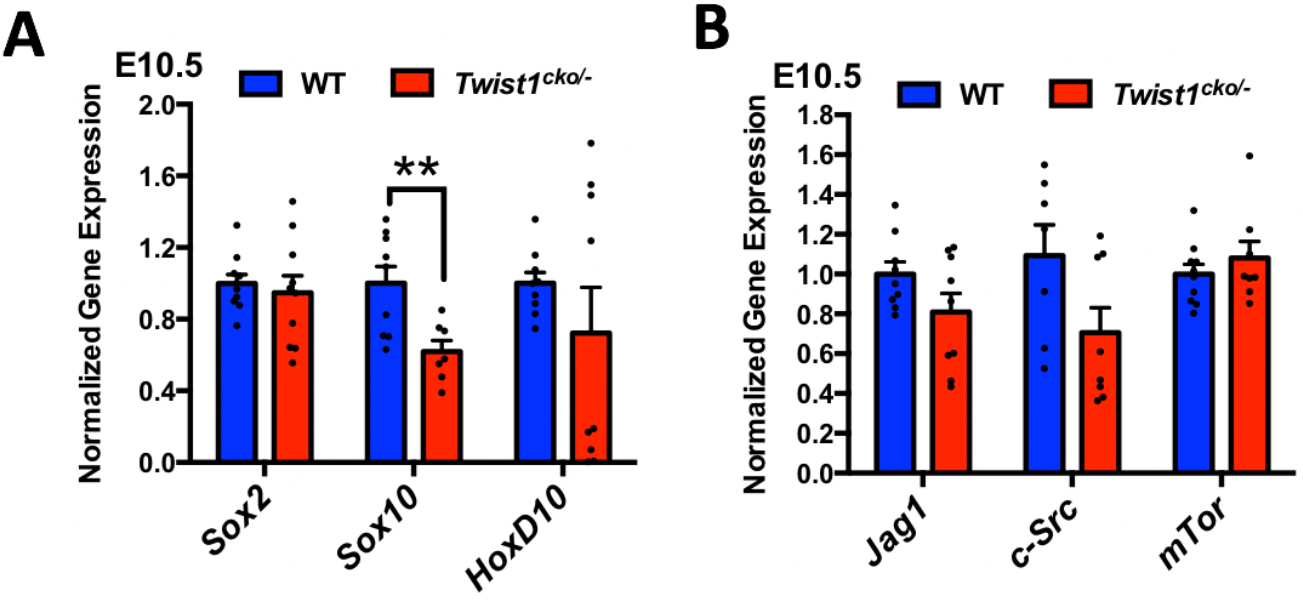
Quantitative measurement of mRNA of transcription factors and signaling ligands in wild type and *Twist1^cko/−^*. (A) mRNA expression of *Sox10* is significantly reduced in *Twist1^cko/−^* tissues compared to WT. (B) mRNA expression of *Jag1*, *c-Src* and *mTor* was not altered in *Twist1^cko/−^* compared to WT.

**Supplementary Fig. 9.**
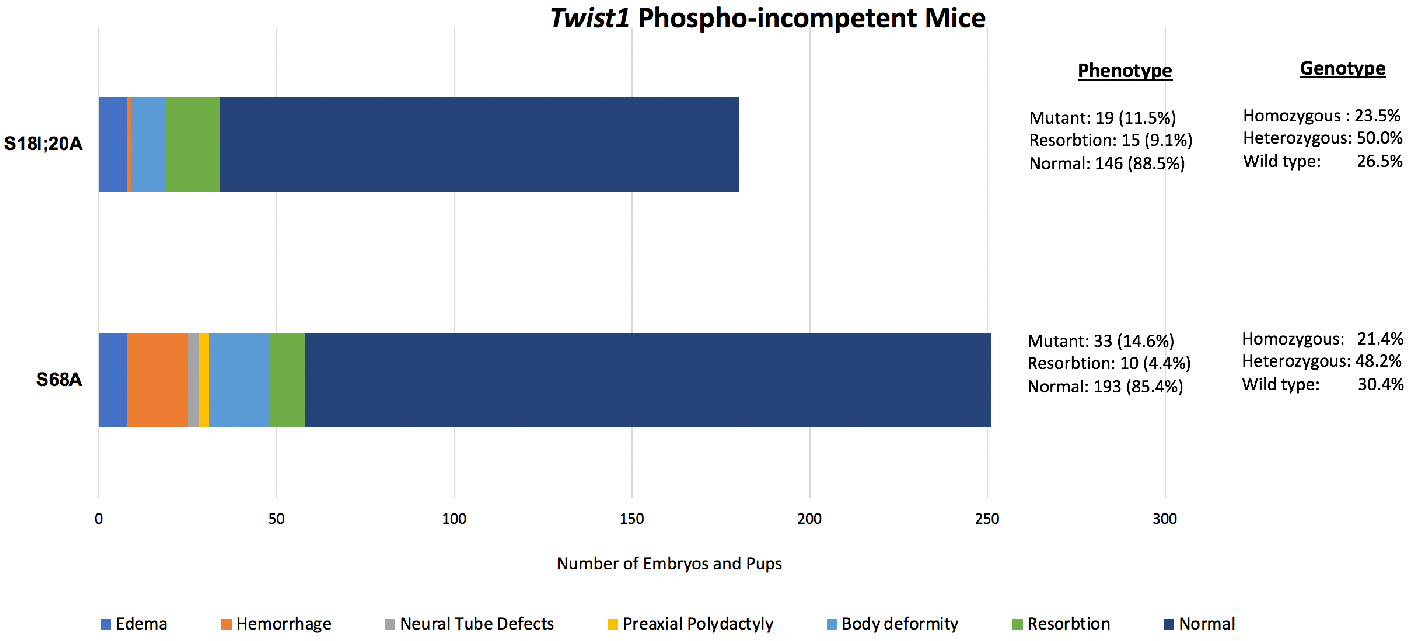
The spectrum of craniofacial phenotype in the homozygous phospho-incompetent *Twist1^S18;20A/S18;20A^* and *Twist1^S68A/S68A^* embryos. The histogram shows the percentage of each phenotype of *Twist1^S18;20A/S18;20A^* and *Twist1^S68A/S68A^* embryos and the percentage of each genotype. The percentage of mutant embryos is less than the estimated genotype percentage of the homozygous phospho-incompetent embryos.

**Supplementary Fig. 10.**
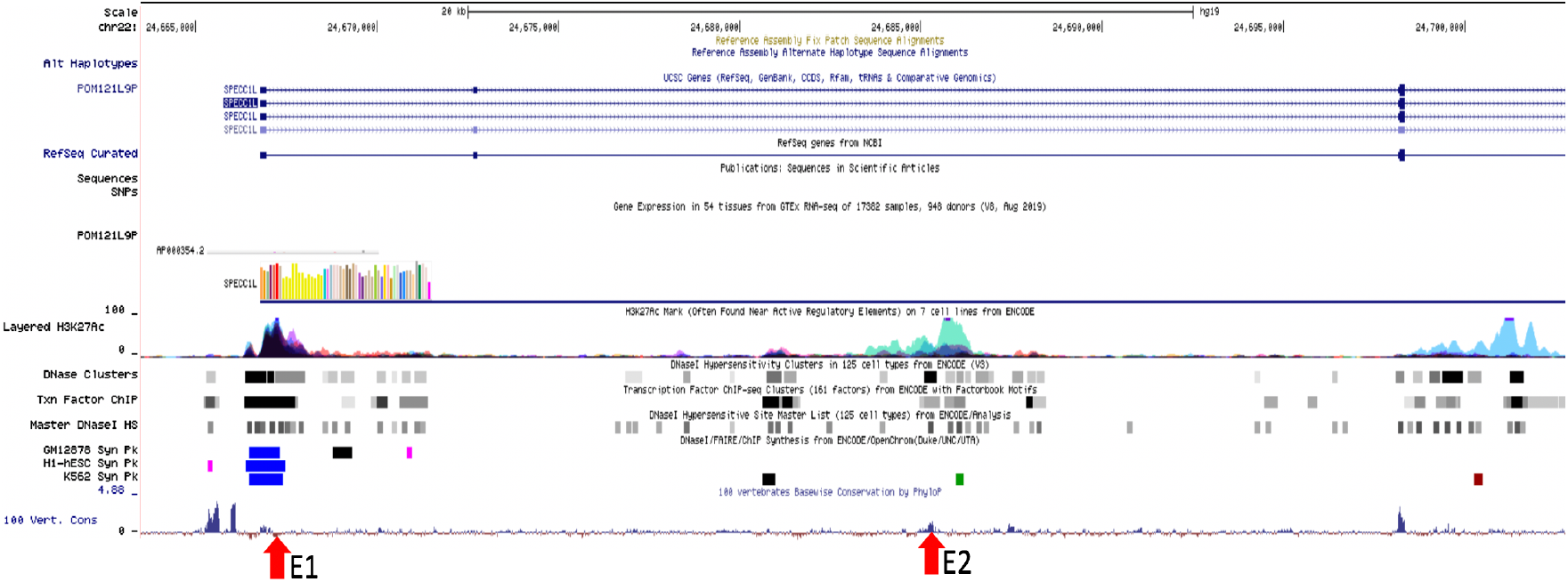
A screenshot from UCSC Genome Browser showing the SPECC1L region. The two red arrows at the bottom indicate the genomic localization of putative enhancer elements E1 and E2 tested for TWIST1 binding. The two regions show enrichment for epigenetic enhancer signatures, including H3K27Ac, DNase I footprint, transcription factor signals, and conservation in 100 vertebrate species.

## References

Achilleos, A., Trainor, P. (2012). Neural crest stem cells: discovery, properties and potential for therapy. Cell Res. 22, 288–304. https://doi.org/10.1038/cr.2012.11

Alexander, N. R., Tran, N. L., Rekapally, H., Summers, C. E., Glackin, C. and Heimark, R.L. (2006). N-cadherin gene expression in prostate carcinoma is modulated by integrin-dependent nuclear translocation of Twist1. Cancer Res. 66, 3365–3369.

Bai, S., Geng, Y., Duan, H., Xu, L., Yu, Z., Yuan, J., & Wei, M. (2021). A novel p.Pro871Leu missense mutation in SPECC1L gene causing craniosynostosis in a patient. Orthodontics and craniofacial research 10.1111/ocr.12473. Advance online publication.

Barrallo-Gimeno, A. and Nieto, M. A. (2005). The Snail genes as inducers of cell movement and survival: implications in development and cancer. Development 132, 3151–3161.

Beck, B., Lapouge, G., Rorive, S., Drogat, B., Desaedelaere, K., Delafaille, S., Dubois, C., Salmon, I., Willekens, K., Marine, J.C. and Blanpain, C. (2015). Different levels of Twist1 regulate skin tumor initiation, stemness, and progression. Cell Stem Cell 16, 67–79.

Bhoj, E. J., Haye, D., Toutain, A., Bonneau, D., Nielsen, I. K., Lund, I. B., Bogaard, P., Leenskjold, S., Karaer, K., Wild, K. T., Grand, K. L., Astiazaran, M. C., Gonzalez-Nieto, L. A., Carvalho, A., Lehalle, D., Amudhavalli, S. M., Repnikova, E., Saunders, C., Thiffault, I., … Verloes, A. (2019). Phenotypic spectrum associated with SPECC1L pathogenic variants: New families and critical review of the nosology of Teebi, Opitz GBBB, and Baraitser-Winter syndromes. European Journal of Medical Genetics, 62(12), 103588.

Bildsoe, H., Fan, X., Wilkie, E. E., Ashoti, A., Jones, V.J., Power, M., Qin, J., Wang, J., Tam, P. P. L. and Loebel, D. A. F. (2016). Transcriptional targets of TWIST1 in the cranial mesoderm regulate cell-matrix interactions and mesenchyme maintenance. Dev. Biol. 418, 189–203.

Bildsoe, H., Loebel, D. A., Jones, V. J., Chen, Y. T., Behringer, R. R. and Tam, P. P. (2009). Requirement for Twist1 in frontonasal and skull vault development in the mouse embryo. Dev. Bio.l 331, 176–188.

Bildsoe, H., Loebel, D. A., Jones, V. J., Hor, A. C., Braithwaite, A. W., Chen, Y. T., Behringer, R. R. and Tam, P. P. (2013). The mesenchymal architecture of the cranial mesoderm of mouse embryos is disrupted by the loss of Twist1 function. Dev. Biol. 374, 295–307.

Bourguignon, L. Y., Wong, G., Earle, C., Krueger, K. and Spevak, C. C. (2010). Hyaluronan-CD44 interaction promotes c-Src-mediated twist signaling, microRNA-10b expression, and RhoA/RhoC up-regulation, leading to Rho-kinase-associated cytoskeleton activation and breast tumor cell invasion. J. Biol. Chem. 285, 36721–36735.

Chai, Y. and Maxson, R. E., Jr. (2006). Recent advances in craniofacial morphogenesis. Dev. Dyn. 235, 2353–2375.

Chang, Y. W., Su, Y. J., Hsiao, M., Wei, K. C., Lin, W. H., Liang, C. L., Chen, S. C. and Lee, J. L. (2015). Diverse Targets of beta-Catenin during the Epithelial-Mesenchymal Transition Define Cancer Stem Cells and Predict Disease Relapse. Cancer Res. 75, 3398–3410.

Chen, J., Yuan, W., Wu, L., Tang, Q., Xia, Q., Ji, J., Liu, Z., Ma, Z., Zhou, Z., Cheng, Y. and Shu, X. (2017). PDGF-D promotes cell growth, aggressiveness, angiogenesis and EMT transformation of colorectal cancer by activation of Notch1/Twist1 pathway. Oncotarget 8, 9961–9973.

Chen, Y. T., Akinwunmi, P. O., Deng, J. M., Tam, O. H. and Behringer, R. R. (2007). Generation of a Twist1 conditional null allele in the mouse. Genesis 45, 588–592.

Chen, Z. F. and Behringer, R. R. (1995). Twist is required in head mesenchyme for cranial neural tube morphogenesis. Genes Dev. 9, 686–699.

Claes, P., Roosenboom, J., White, J. D., Swigut, T., Sero, D., Li, J., Lee, M. K., Zaidi, A., Mattern, B. C., Liebowitz, C., Pearson, L., González, T., Leslie, E. J., Carlson, J. C., Orlova, E., Suetens, P., Vandermeulen, D., Feingold, E., Marazita, M. L., Shaffer, J. R., … Weinberg, S. M. (2018). Genome-wide mapping of global-to-local genetic effects on human facial shape. Nature Genetics 50 (3), 414–423.

Cousin, H. (2017) Cadherins function during the collective cell migration of Xenopus Cranial Neural Crest cells: revisiting the role of E-cadherin. Mech Dev. 148:79–88. doi: 10.1016/j.mod.2017.04.006. Epub 2017 Apr 30. PMID: 28467887; PMCID: PMC5662486.

Eckert, M. A., Lwin, T. M., Chang, A. T., Kim, J., Danis, E., Ohno-Machado, L. and Yang, J. (2011). Twist1-induced invadopodia formation promotes tumor metastasis. Cancer Cell 19, 372–386.

Fakhouri, W. D., Metwalli, K., Naji, A., Bakhiet, S., Quispe-Salcedo, A., Nitschke, L., Kousa, Y. A., and Schutte, B. C. (2017). Intercellular Genetic Interaction Between *Irf6* and *Twist1* during Craniofacial Development. Sci. Rep. 7, 7129.

Fakhouri, W. D., Rhea, L., Du, T., Sweezer, E., Morrison, H., Fitzpatrick, D., Yang, B., Dunnwald, M. and Schutte, B. C. (2012). MCS9.7 enhancer activity is highly, but not completely, associated with expression of Irf6 and p63. Dev. Dyn. 241, 340–349.

Fan, X., Masamsetti, V. P., Sun, J. Q., Engholm-Keller, K., Osteil, P., Studdert, J., Graham, M. E., Fossat, N., Tam, P. P. (2021). TWIST1 and chromatin regulatory proteins interact to guide neural crest cell differentiation. Elife. 10:e62873. doi: 10.7554/eLife.62873. PMID: 33554859; PMCID: PMC7968925.

Farrell, B. C., Power, E. M., Mc Dermott, K. W. (2011). Developmentally regulated expression of Sox9 and microRNAs 124, 128 and 23 in neuroepithelial stem cells in the developing spinal cord. Int J Dev Neurosci. 29(1):31–6. doi: 10.1016/j.ijdevneu.2010.10.001. Epub 2010 Oct 16. PMID: 20937378.

Firulli, A. B. and Conway, S. J. (2008). Phosphoregulation of Twist1 provides a mechanism of cell fate control. Curr. Med. Chem. 15, 2641–2647.

Firulli, B. A., Krawchuk, D., Centonze, V. E., Vargesson, N., Virshup, D. M., Conway, S. J., Cserjesi, P., Laufer, E. and Firulli, A. B. (2005). Altered Twist1 and Hand2 dimerization is associated with Saethre-Chotzen syndrome and limb abnormalities. Nat. Genet. 37, 373–381.

Firulli, B. A., Redick, B. A., Conway, S. J., Firulli, A. B. (2007) Mutations within helix I of Twist1 result in distinct limb defects and variation of DNA binding affinities. J Biol Chem. 282(37):27536–27546. doi: 10.1074/jbc.M702613200. Epub 2007 Jul 25. PMID: 17652084; PMCID: PMC2556885

Füchtbauer, E. M. (1995). Expression of M-twist during postimplantation development of the mouse. Dev Dyn. 204(3):316–22. doi: 10.1002/aja.1002040309. PMID: 8573722.

Gfrerer, L., Shubinets, V., Hoyos, T., Kong, Y., Nguyen, C., Pietschmann, P., Morton, C. C., Maas, R. L., Liao, E. C. (2014). Functional analysis of SPECC1L in craniofacial development and oblique facial cleft pathogenesis. Plastic and reconstructive surgery, 134(4), 748–759.

Goering, J. P., Isai, D. G., Hall, E. G., Wilson, N. R., Kosa, E., Wenger, L. W., Umar, Z., Yousaf, A., Czirok, A., Saadi, I. (2021). SPECC1L-deficient primary mouse embryonic palatal mesenchyme cells show speed and directionality defects. Scientific Reports, 11.

Groves, A. K., LaBonne, C. (2014). Setting appropriate boundaries: fate, patterning and competence at the neural plate border. Dev Biol. 1;389(1):2–12. doi: 10.1016/j.ydbio.2013.11.027. Epub 2013 Dec 7. PMID: 24321819; PMCID: PMC3972267.

He, F., Hu, X., Xiong, W., Li, L., Lin, L., Shen, B., Yang, L., Gu, S., Zhang, Y. and Chen, Y. (2014). Directed Bmp4 expression in neural crest cells generates a genetic model for the rare human bony syngnathia birth defect. Dev. Biol. 391, 170–181.

Hong, J., Zhou, J., Fu, J., He, T., Qin, J., Wang, L., Liao, L. and Xu, J. (2011). Phosphorylation of serine 68 of Twist1 by MAPKs stabilizes Twist1 protein and promotes breast cancer cell invasiveness. Cancer Res. 71, 3980–3990.

Hoyert, D. L., Mathews, T. J., Menacker, F., Strobino, D. M. and Guyer, B. (2006). Annual summary of vital statistics: 2004. Pediatrics 117, 168–183.

Ingraham, C. R., Kinoshita, A., Kondo, S., Yang, B., Sajan, S., Trout, K. J., Malik, M. I., Dunnwald, M., Goudy, S. L., Lovett, M., Murray, J. C. and Schutte, B. C. (2006). Abnormal skin, limb and craniofacial morphogenesis in mice deficient for interferon regulatory factor 6 (Irf6). Nat. Genet. 38, 1335–1340.

Joshi, N., Hamdan, A. M. and Fakhouri, W. D. (2014). Skeletal malocclusion: a developmental disorder with a life-long morbidity. J. Clin. Med. Res. 6, 399–408.

Kousa, Y. A., Zhu, H., Fakhouri, W. D., Lei, Y., Kinoshita, A., Roushangar, R. R., Patel, N. K., Agopian, A. J., Yang, W., Leslie, E. J., et al. (2019). The TFAP2A-IRF6-GRHL3 genetic pathway is conserved in neurulation. Hum. Mol. Genet. 28, 1726–1737.

Kunz, J., Hudler, M. and Fritz, B. (1999). Identification of a frameshift mutation in the gene TWIST in a family affected with Robinow-Sorauf syndrome. J. Med. Genet. 36, 650–652.

Le Douarin, N. M., Creuzet, S., Couly, G. and Dupin, E. (2004). Neural crest cell plasticity and its limits. Development 131, 4637–4650.

Lewis, A. E., Vasudevan, H. N., O’Neill, A. K., Soriano, P., Bush, J. O. (2013). The widely used Wnt1-Cre transgene causes developmental phenotypes by ectopic activation of Wnt signaling. Developmental biology, 379 (2), 229–234. https://doi.org/10.1016/j.ydbio.2013.04.026

Litingtung, Y., Chiang, C. (2000). Specification of ventral neuron types is mediated by an antagonistic interaction between Shh and Gli3. Nat. Neurosci. 3, 979–985.

Lu, S., Nie, J., Luan, Q., Feng, Q., Xiao, Q., Chang, Z., Shan, C., Hess, D., Hemmings, B. A. and Yang, Z. (2011). Phosphorylation of the Twist1-family basic helix-loop-helix transcription factors is involved in pathological cardiac remodeling. PLoS One 6, e19251.

Metwalli, K. A., Do, M. A., Nguyen, K., Mallick, S., Kin, K., Farokhnia, N., Jun, G., Fakhouri, W. D. (2018). Interferon Regulatory Factor 6 Is Necessary for Salivary Glands and Pancreas Development. Journal of dental research 97(2), 226–236.

Millonig, J. H., Millen, K. J., Hatten, M. E. (2000) The mouse Dreher gene Lmx1a controls formation of the roof plate in the vertebrate CNS. Nature 403, 764–769.

Murray, S. A., Oram, K. F. and Gridley, T. (2007). Multiple functions of Snail family genes during palate development in mice. Development 134, 1789–1797.

Nieto, M. A., Huang, R. Y., Jackson, R. A. and Thiery, J. P. (2016). Emt: 2016. Cell 166, 21–45.

Noden, D. M. and Trainor, P. A. (2005). Relations and interactions between cranial mesoderm and neural crest populations. J. Anat. 207, 575–601.

Nomura, M. and Li, E. (1998). Smad2 role in mesoderm formation, left-right patterning and craniofacial development. Nature 393, 786–790.

Nusslein-Volhard, C., Wieschaus, E., Kluding, H. (1984). Mutations affecting the pattern of the larval cuticle in Drosophila melanogaster. 1. Zygotic loci on the 2nd chromosome. Roux’s Arch. Dev. Biol. 193, 267–282.

Ota, M. S., Loebel, D. A., O’Rourke, M. P., Wong, N., Tsoi, B., & Tam, P. P. (2004). Twist is required for patterning the cranial nerves and maintaining the viability of mesodermal cells. Developmental dynamics : an official publication of the American Association of Anatomists, 230(2), 216–228. https://doi.org/10.1002/dvdy.20047

Padmanabhan, R. and Ahmed, I. (1997). Retinoic acid-induced asymmetric craniofacial growth and cleft palate in the TO mouse fetus. Reprod. Toxicol. 11, 843–860.

Parker, S. E., Mai, C. T., Canfield, M. A., Rickard, R., Wang, Y., Meyer, R. E., Anderson, P., Mason, C. A., Collins, J. S., Kirby, R. S. and Correa, A. (2010). Updated National Birth Prevalence estimates for selected birth defects in the United States, 2004-2006. Birth Defects Res. A. Clin. Mol. Teratol 88, 1008–1016.

Parr, B. A., Shea, M. J., Vassileva, G. and McMahon, A. P. (1993). Mouse Wnt genes exhibit discrete domains of expression in the early embryonic CNS and limb buds. Development 119, 247–261.

Peyrard-Janvid, M., Leslie, E. J., Kousa, Y. A., Smith, T. L., Dunnwald, M., Magnusson, M., Lentz, B. A., Unneberg, P., Fransson, I., Koillinen, H. K., et al (2014). Dominant mutations in GRHL3 cause Van der Woude Syndrome and disrupt oral periderm development. Am. J. Hum. Genet. 94, 23–32.

Pieters, T., Sanders, E., Tian, H., van Hengel, J., van Roy, F. (2020). Neural defects caused by total and Wnt1-Cre mediated ablation of p120ctn in mice. BMC developmental biology, 20 (1), 17.

Richardson, R. J., Dixon, J., Malhotra, S., Hardman, M. J., Knowles, L., Boot-Handford, R. P., Shore, P., Whitmarsh and A., Dixon, M. J. (2006). Irf6 is a key determinant of the keratinocyte proliferation-differentiation switch. Nat. Genet. 38, 1329–1334.

Rogers, C. D., Saxena, A. and Bronner, M. E. (2013). Sip1 mediates an E-cadherin-to-N-cadherin switch during cranial neural crest EMT. J Cell Biol 203, 835–847.

Saadi, I., Alkuraya, F. S., Gisselbrecht, S. S., Goessling, W., Cavallesco, R., Turbe-Doan, A., Petrin, A. L., Harris, J., Siddiqui, U., Grix, A. W., Hove, H. D., Leboulch, P., Glover, T. W., Morton, C. C., Richieri-Costa, A., Murray, J. C., Erickson, R. P., Maas, R. L. (2011). Deficiency of the Cytoskeletal Protein SPECC1L Leads to Oblique Facial Clefting. American Journal of Human Genetics, 89 (1), 44–55.

Sato, T., Kurihara, Y., Asai, R., Kawamura, Y., Tonami, K., Uchijima, Y., Heude, E., Ekker, M., Levi, G. and Kurihara, H. (2008). An endothelin-1 switch specifies maxillomandibular identity. Proc. Natl. Acad. Sci. U. S. A. 105, 18806–18811.

Seto, M. L., Hing, A. V., Chang, J., Hu, M., Kapp-Simon, K. A., Patel, P. K., Burton, B. K., Kane, A. A., Smyth, M. D., Hopper, R., et al. (2007). Isolated sagittal and coronal craniosynostosis associated with TWIST box mutations. Am. J. Med. Genet. A. 143A, 678–686.

Simpson, P. (1983). Maternal–zygotic gene interactions during formation of the dorsoventral pattern in Drosophila embryos. Genetics 105, 615–632.

Soldatov, R., Kaucka, M., Kastriti, M. E., Petersen, J., Chontorotzea, T., Englmaier, L., Akkuratova, N., Yang, Y., Haring, M., Dyachuk, V., et al. (2019). Spatiotemporal structure of cell fate decisions in murine neural crest. Science 364.

Soo K, O’Rourke MP, Khoo PL, Steiner KA, Wong N, Behringer RR, Tam PP. (2002). Twist function is required for the morphogenesis of the cephalic neural tube and the differentiation of the cranial neural crest cells in the mouse embryo. Dev Biol. 247(2):251–70. doi: 10.1006/dbio.2002.0699. PMID: 12086465.

Stoetzel, C., Weber, B., Bourgeois, P., Bolcato-Bellemin, A. L., Perrin-Schmitt, F. Dorso-ventral and rostro-caudal sequential expression of M-twist in the postimplantation murine embryo. Mechanisms of development, 1995 51 (2-3), 251–263.

Su, Y. W., Xie, T. X., Sano, D. and Myers, J. N. (2011). IL-6 stabilizes Twist and enhances tumor cell motility in head and neck cancer cells through activation of casein kinase 2. PLoS One 6, e19412.

Takenouchi, T., Sakamoto, Y., Sato, H., Suzuki, H., Uehara, T., Ohsone, Y. and Kosaki, K. (2018). Ablepharon and craniosynostosis in a patient with a localized TWIST1 basic domain substitution. Am. J. Med. Genet. A. 176, 2777–2780.

Thiery, J. P., Acloque, H., Huang, R. Y. and Nieto, M. A. (2009). Epithelial-mesenchymal transitions in development and disease. Cell 139, 871–890.

Thompson, J., Mendoza, F., Tan, E., Bertol, J. W., Gaggar, A. S., Jun, G., Biguetti, C. and Fakhouri, W. D. (2019). A cleft lip and palate gene, Irf6, is involved in osteoblast differentiation of craniofacial bone. Dev. Dyn. 248, 221–232.

Vandewalle, C., Comijn, J., De Craene, B., Vermassen, P., Bruyneel, E., Andersen, H., Tulchinsky, E., Van Roy, F. and Berx, G. (2005). SIP1/ZEB2 induces EMT by repressing genes of different epithelial cell-cell junctions. Nucleic Acids Res. 33, 6566–6578.

Vesuna, F., van Diest, P., Chen, J. H., Raman, V. (2008). Twist is a transcriptional repressor of E-cadherin gene expression in breast cancer. Biochemical and biophysical research communications, 367 (2), 235–241.

Vichalkovski, A., Gresko, E., Hess, D., Restuccia, D. F. and Hemmings, B. A. (2010). PKB/AKT phosphorylation of the transcription factor Twist-1 at Ser42 inhibits p53 activity in response to DNA damage. Oncogene 29, 3554–3565.

Vincentz, J. W., Barnes, R. M., Rodgers, R., Firulli, B. A., Conway, S. J. and Firulli, A. B. (2008). An absence of Twist1 results in aberrant cardiac neural crest morphogenesis. Dev. Biol. 320, 131–139.

Vogel, J. K., Wegner, M. (2021). Sox9 in the developing central nervous system: a jack of all trades? Neural Regen Res. 16 (4):676–677. doi: 10.4103/1673-5374.295327. PMID: 33063721; PMCID: PMC8067939.

Wang, Y., Shi, J., Chai, K., Ying, X. and Zhou, B.P. (2013). The Role of Snail in EMT and Tumorigenesis. Curr. Cancer Drug Targets 13, 963–972

Whipple, R. A., Matrone, M. A., Cho, E. H., Balzer, E. M., Vitolo, M. I., Yoon, J. R., Ioffe, O. B., Tuttle, K. C., Yang, J., Martin, S. S. (2010). Epithelial-to-mesenchymal transition promotes tubulin detyrosination and microtentacles that enhance endothelial engagement. Cancer Research, 70 (20), 8127–8137.

White, J. D., Indencleef, K., Naqvi, S., Eller, R. J., Hoskens, H., Roosenboom, J., Lee, M. K., Li, J., Mohammed, J., Richmond, S., Quillen, E. E., Norton, H. L., Feingold, E., Swigut, T., Marazita, M. L., Peeters, H., Hens, G., Shaffer, J. R., Wysocka, J., Walsh, S., … Claes, P. (2021). Insights into the genetic architecture of the human face. Nature Genetics, 53 (1), 45–53.

Wilson, N. R., Olm-Shipman, A. J., Acevedo, D. S., Palaniyandi, K., Hall, E. G., Kosa, E., Stumpff, K. M., Smith, G. J., Pitstick, L., Liao, E. C., Bjork, B. C., Czirok, A., Saadi, I. (2016). SPECC1L deficiency results in increased adherens junction stability and reduced cranial neural crest cell delamination. Scientific Reports, 6.

Wolf, C., Thisse, C., Stoetzel, C., Thisse, B., Gerlinger, P., Perrin-Schmitt, F. (1991). The M-twist gene of Mus is expressed in subsets of mesodermal cells and is closely related to the Xenopus X-twi and the Drosophila twist genes. Developmental biology, 143 (2), 363–373.

Xu, Y., Lee, D. K., Feng, Z., Xu, Y., Bu, W., Li, Y., Liao, L., Xu, J. (2017). Breast tumor cell-specific knockout of Twist1 inhibits cancer cell plasticity, dissemination, and lung metastasis in mice. Proc. Natl. Acad. Sci. U. S. A. 114, 11494–11499.

Xue, G. and Hemmings, B. A. (2012). Phosphorylation of basic helix-loop-helix transcription factor Twist in development and disease. Biochem Soc. Trans. 40, 90–93.

Zhang, A., Aslam, H., Sharma, N., Warmflash, A., Fakhouri, W. D. (2021). Conservation of Epithelial-to-Mesenchymal Transition Process in Neural Crest Cells and Metastatic Cancer. Cells, Tissues, Organs, 1–22. Advance online publication. https://doi.org/10.1159/000516466

Zhang, Y., Blackwell, E. L., McKnight, M. T., Knutsen, G. R., Vu, W. T. and Ruest, L. B. (2012). Specific inactivation of Twist1 in the mandibular arch neural crest cells affects the development of the ramus and reveals interactions with hand2. Dev. Dyn. 241, 924–940.

Zhao, Z., Rahman, M. A., Chen, Z. G., Shin, D. M. (2017). Multiple biological functions of Twist1 in various cancers. Oncotarget, 8 (12), 20380–20393.

